# High-resolution measurement of ERK activation state reveals determinants of BRAF regulation and drug sensitivity

**DOI:** 10.64898/2026.09.30.755370

**Authors:** A. Felicia Adebanjo, Jessica J. Simon, Fernando Banales-Mejia, Douglas M. Fowler, Dustin J. Maly

**Affiliations:** Department of Chemistry, University of Washington, Seattle, WA 98195, USA; Department of Genome Sciences, University of Washington, Seattle, WA 98195, USA; Department of Bioengineering, University of Washington, Seattle, WA 98195, USA; Department of Biochemistry, University of Washington, Seattle, WA 98195, USA

## Abstract

High-resolution measurements of pathway activity make it possible to characterize how signaling responds to changing cellular conditions, to evaluate pathway-targeted inhibitors, and to dissect how sequence variation shapes signaling output. Here, we describe a sequencing-based assay that uses engineered, phospho-state-selective affinity reagents to quantify ERK activation state as a high-resolution, ratiometric readout of RAS/ERK MAPK pathway activity. Profiling ∼1,600 BRAF variants at basal and elevated RAS-GTP levels allowed us to assign activity scores to somatic cancer and germline RASopathy variants and to distinguish residues that stabilize autoinhibition from those required for active dimer formation. We identified a large class of dominant negative variants that suppress signaling below endogenous levels through enhanced RAS engagement and an inability to form productive RAF dimers. Measuring the intracellular potency of the RAF inhibitor LXH254 against ∼1,000 variants in parallel revealed hinge and activation-loop positions conferring resistance and showed that disrupting the BRAF-MEK interface is broadly sensitizing.

## Introduction

Cell behavior and fate are governed by intracellular signaling pathways that convert extracellular and intracellular cues into appropriate cellular responses, ranging from survival and proliferation to apoptosis and metabolic adaptation.^1^ The activity of these pathways is highly dynamic, continuously tuned by the cellular environment, including growth factor availability, the activation state of upstream receptors, and feedback from downstream effectors.^2,3^ Assays that provide high-resolution measurements of pathway activity are therefore valuable, revealing how a pathway responds to changes in cellular conditions, such as growth factor stimulation or the expression of an oncogene. They also allow the high-precision evaluation of how pathway-targeted inhibitors affect pathway output. Assays that can be adapted to the profiling of protein variant panels and libraries of proteins are especially powerful, as they enable systematic analysis of how individual pathway components shape pathway output and provide insight into both basic signaling mechanisms and the functional consequences of disease-associated mutations.^4,5^

The RAS/ERK mitogen-activated protein kinase (MAPK) pathway is among the most extensively studied signaling cascades, controlling cell proliferation, differentiation, survival, and migration across diverse cell types.^6^ Tools for quantifying the activity of this pathway are particularly valuable because of the breadth of the cellular processes it governs and the central role of its dysregulation in cancer. Genes encoding proteins within this pathway, including RAS and RAF family members, are among the most frequently mutated oncogenes,^7^ and many of the proteins that regulate MAPK pathway activity are themselves established drug targets.^8^ ERK, the terminal kinase of the cascade, is an especially informative readout of pathway activity because its activation state integrates input from numerous upstream proteins: cell-surface receptors of several classes, including receptor tyrosine kinases (RTKs), integrins, and G protein-coupled receptors (GPCRs), RAS GTPases, guanine nucleotide exchange factors (GEFs), GTPase-activating proteins (GAPs), adaptor proteins, and RAF and MEK kinases.^9,10^ As a result, assays that use ERK activation state as a readout can measure the functional consequences of perturbing any of these components.

Here, we describe the development and application of a high-resolution, next-generation-sequencing (NGS)-compatible assay for quantifying ERK activation state and, consequently, RAS/ERK MAPK pathway activity. The assay relies on optimized sequential enrichment of the unphosphorylated and phosphorylated forms of an ERK reporter bound to a circular RNA barcode, providing a ratiometric, NGS-based measurement of ERK activation state. We used this assay to systematically profile a library of ∼1,600 BRAF variants at basal and elevated RAS-GTP levels, enabling us to assign quantitative pathway activity scores to clinically observed somatic and germline variants and to dissect the structural basis of BRAF autoinhibition and active dimer formation. We also identified a diversity of dominant negative variants that suppress pathway activity below endogenous levels and characterized variant-specific responses to the RAF inhibitor LXH254 (naporafenib). Together, these results demonstrate the utility of high-resolution, sequencing-based pathway activity measurements for resolving sequence-function relationships in signaling proteins and provide further mechanistic insight into the role of BRAF in the MAPK pathway.

## Results

### DARPin-based enrichment improves resolution and dynamic range of ERK activation state measurements

ERK is the terminal kinase of the RAS/ERK MAPK signaling pathway, and its activation state, which is directly reflected by the phosphorylation level of its activation loop, can be used as a readout of pathway activity (**Figure 1A**). We previously developed an assay based on the LABEL-seq platform for quantifying ERK activation state. This assay requires exogenous expression of a FLAG-tagged reporter in which a tandem dimer of the RNA-binding protein MCP (tdMCP) is fused to the kinase ERK2. Inside cells, the tdMCP moiety assembles with a coexpressed circular RNA (circRNA) bearing an identifying barcode.^5^ Overall pathway activity was measured by selectively immunoprecipitating activation loop-phosphorylated tdMCP-ERK2 with a phospho-ERK(pERK) -specific antibody after cell lysis, followed by reverse transcription and NGS-based barcode quantification normalized to total barcode frequency (**Figure 1B**). We used this assay to quantify ERK activation driven by a pooled library of ∼1,600 BRAF variants with variable capacities to promote downstream signaling, and found that pathway activity scores correlated well with pERK levels measured by western blot and were highly reproducible.^5^ However, the assay did not achieve the dynamic range of western blot analysis.

**Figure 1:**
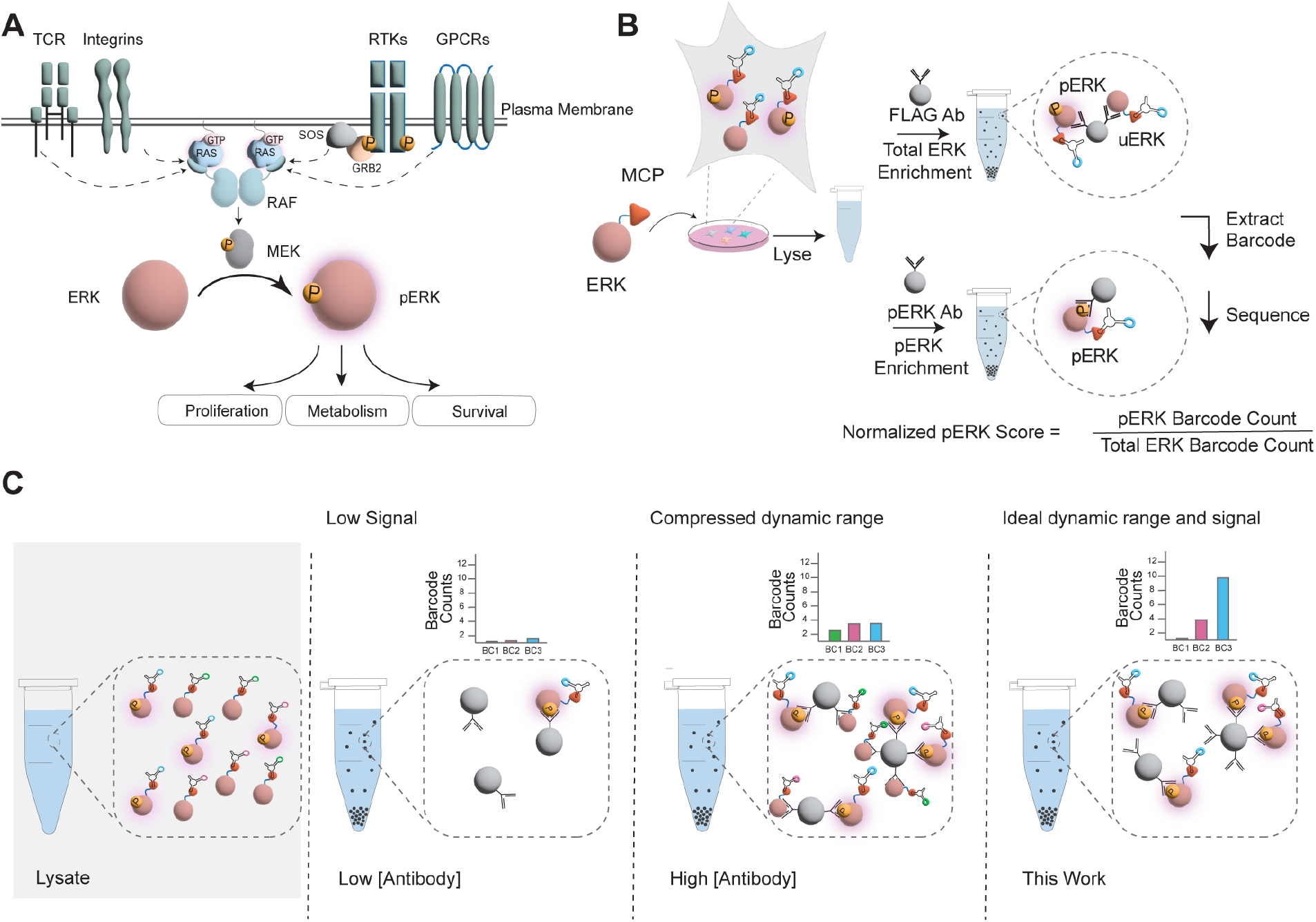
A sequencing-based ERK activation-state reporter for measuring MAPK pathway activity. **(A)** Schematic of the RAS/ERK mitogen-activated protein kinase (MAPK) signaling pathway and the cell-surface receptors that control its activation state, including T cell receptors (TCRs), integrins, receptor tyrosine kinases (RTKs), and G protein-coupled receptors (GPCRs). **(B)** Workflow of the previously reported LABEL-seq pathway activity assay, which used antibody-based enrichment to measure ERK activation state.^5^ Cells stably expressing a barcoded circular RNA (circRNA) also express a 3xFLAG-tagged tdMCP-ERK2 fusion reporter, which assembles intracellularly with the circRNA. Following lysis, phosphorylated ERK (pERK) is enriched with an anti-pERK antibody, and total reporter is enriched in parallel with an anti-FLAG antibody for normalization. Barcodes from each enrichment are extracted and quantified by next-generation sequencing (NGS), and pathway activity is calculated as the pERK/FLAG barcode ratio, normalized to WT. **(C)** Conceptual illustration of the trade-off between signal and specificity in pERK enrichment. Low concentrations of the enrichment reagent yield insufficient capture of pERK and its associated circRNA barcodes, whereas high concentrations promote nonspecific co-capture of unphosphorylated ERK (uERK). An ideal enrichment strategy provides efficient capture of pERK with minimal co-capture of uERK.

We hypothesized that the reduced dynamic range of our pathway activity assay resulted from the high antibody concentrations required to capture the majority of phosphorylated tdMCP-ERK2 prior to barcode quantification, which increases the likelihood of co-enrichment of the lower-affinity unphosphorylated form (**Figure 1C**). This problem is particularly acute for phosphorylated signaling enzymes, where the unphosphorylated species are often present at substantially higher concentrations than their phosphorylated counterparts.^11^ To overcome this limitation, we explored the use of designed ankyrin repeat proteins (DARPins) as antibody replacements.^12^ DARPins are monomeric, homogeneous affinity reagents. DARPins selective for both phosphorylated ERK (pERK) and unphosphorylated ERK (uERK) have been developed.^13,14^ We reasoned that precisely controlled concentrations of immobilized DARPin would enable efficient enrichment of the desired ERK species while minimizing co-enrichment of the lower-affinity form.

To implement our DARPin-based enrichment strategy, we generated biotinylated versions of both pERK-selective (pE59 and pEM1) and uERK-selective (E40) DARPins. Using the reported affinities of each DARPin for pERK and uERK, we modeled the fraction of each ERK species bound across a range of DARPin concentrations and identified immobilization concentrations for pE59, pEM1, and E40 predicted to efficiently capture the preferred ERK species with minimal co-enrichment of the non-preferred species (**Figure 2A**). To profile the efficiency and selectivity of each immobilized DARPin at the selected concentrations, we measured enrichment of the GST-tagged preferred ERK species from an equimolar mixture with the non-GST-tagged non-preferred ERK species (**Figure 2B**). We found that E40 provided highly selective and near-quantitative enrichment of GST-uERK, and undetectable co-enrichment of non-GST-tagged pERK across all concentrations tested. Immobilized pE59 and pEM1 demonstrated lower selectivity for pERK over uERK, but we identified concentrations of each that efficiently and near-quantitatively enriched GST-pERK with minimal co-enrichment of non-GST-tagged uERK (**Figure 2B**).

**Figure 2:**
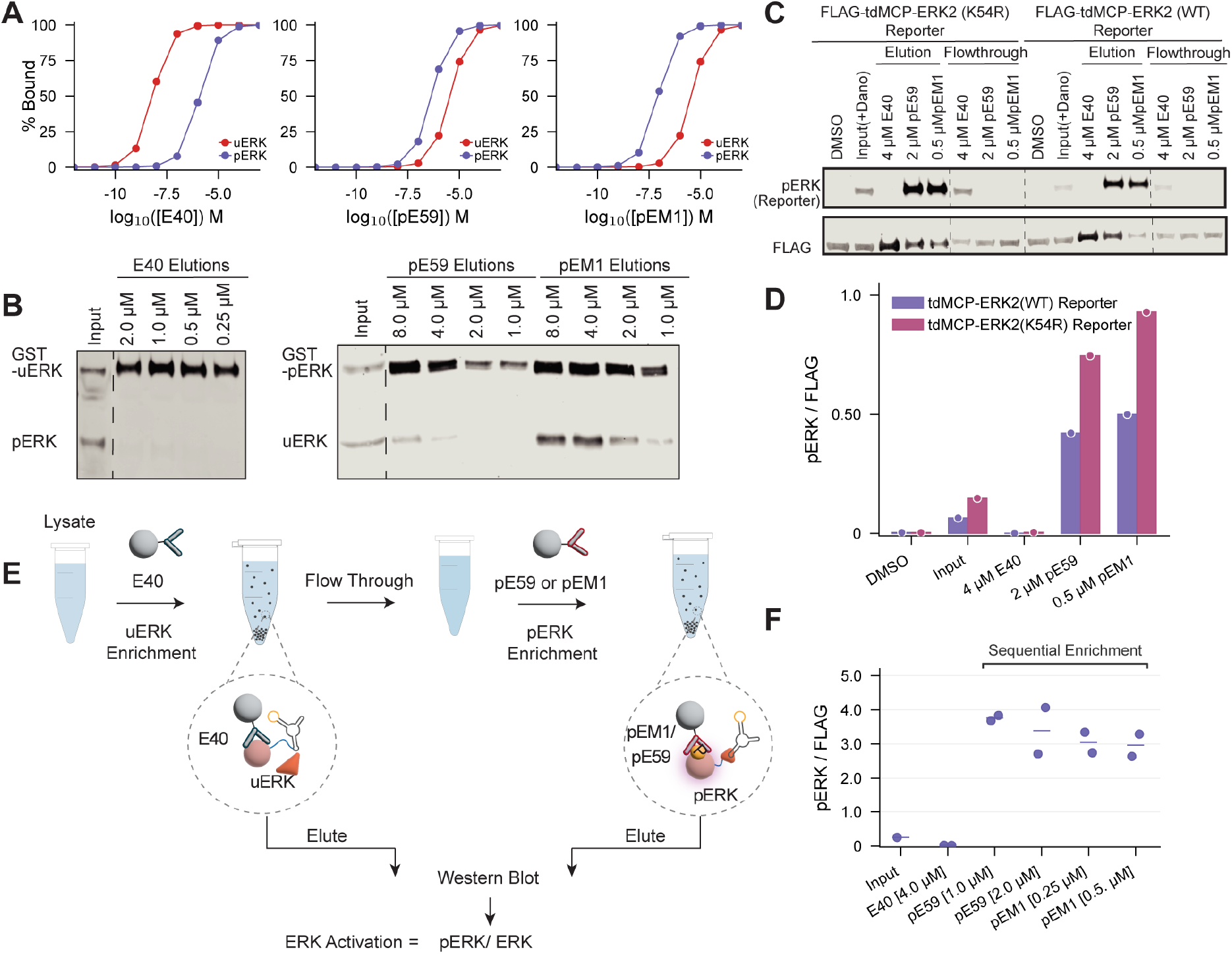
Phospho-state-selective DARPins enable sequential enrichment of a kinase-dead ERK reporter. **(A)** Predicted fractional occupancy of uERK and pERK by the DARPins E40, pE59, and pEM1 across the concentration range shown. Curves give percent bound for uERK (red) and pERK (purple), calculated from reported dissociation constant (K_D_) values.^13,14^ **(B)** Left, western blot showing GST-uERK and untagged pERK enriched from an equimolar mixture across immobilized E40 concentrations of 0.25-2 µM. Right, western blot showing GST-pERK and untagged uERK enriched from the equivalent mixture across immobilized pE59 or pEM1 concentrations of 1-8 µM. In each assay the preferred ERK species carries the GST tag. Input and elution fractions, run on the same gel, are shown for each enrichment (n = 1). **(C)** Western blot of parallel uERK and pERK enrichment from lysates of NS3a-CIAR cells (Figure S1A) transiently expressing WT or kinase-dead (K54R) tdMCP-ERK2 reporters, using immobilized E40, pE59, or pEM1 at the concentrations shown. Before lysis, cells were treated with danoprevir (1 µM, 1 h) to elevate RAS-GTP and raise pERK levels; the leftmost lane of each blot is lysate from DMSO-treated cells, giving a baseline for the phosphorylation level of each reporter under basal conditions (n = 1). **(D)** Quantification of tdMCP-pERK2 for each reporter normalized to total reporter (pERK/FLAG) from the blots in (C) (n = 1). **(E)** Schematic of sequential DARPin-based enrichment. Lysate is first depleted of tdMCP-uERK2 with E40-conjugated streptavidin beads; tdMCP-pERK2 remaining in the flow-through is then enriched with immobilized pE59 or pEM1. Eluted fractions from the two steps are resolved in separate lanes. **(F)** Quantification of FLAG-normalized pERK signal from sequential enrichment of the tdMCP-ERK2(K54R) reporter at the pE59 and pEM1 concentrations indicated, from the blots in Figure S1B. Each dot represents an individual sequential enrichment; bars indicate the mean (n = 2 biological replicates). *See also Figure S1*.

### A kinase-dead ERK reporter supports sequential DARPin-based enrichment of phosphorylated and unphosphorylated species

Having established optimal DARPin concentrations for selective ERK enrichment, we next evaluated whether the tdMCP-ERK2 reporter could be improved prior to implementing the full LABEL-seq workflow. Our previous tdMCP-ERK2 reporter used a catalytically active version of the kinase, which raises the possibility that reporter activation may perturb the cellular signaling state. Therefore, we determined whether the activation loop of a kinase-dead version of ERK (K54R)^15^ could be phosphorylated to levels similar to the catalytically competent version in the context of a tdMCP fusion, and whether immobilized DARPins retained their affinities and selectivities for their preferred phosphorylated and unphosphorylated forms of a tdMCP-ERK2(K54R) reporter. To test this, we compared the phosphorylation levels and DARPin enrichment of tdMCP-ERK2(K54R) and tdMCP-ERK2(WT) reporters in HEK293 cells expressing NS3a-CIAR, which enables temporally controlled increases in endogenous RAS-GTP levels and downstream pathway activation upon treatment with the bioorthogonal small molecule danoprevir^16,17^ (**Figure S1A**). Upon NS3a-CIAR activation, tdMCP-ERK2(K54R) was phosphorylated to slightly higher levels than tdMCP-ERK2(WT), and pE59 and pEM1 quantitatively enriched the phosphorylated form of both reporters (**Figure 2C** and **2D**). Importantly, as with tdMCP-ERK2(WT), E40 enriched negligible levels of phosphorylated tdMCP-ERK2(K54R), demonstrating that the DARPins maintain their affinities and selectivities for both phosphorylated and unphosphorylated forms of the kinase-dead reporter.

The availability of DARPins selective for both uERK and pERK enables a sequential enrichment strategy in which one ERK species is captured and quantified, and the other is subsequently captured and quantified from the flow-through. The ratio of each ERK species captured in each step can then be used to calculate overall ERK activation levels (**Figure 2E**). We reasoned that the high selectivity of E40 for uERK, combined with the likely higher abundance of uERK relative to pERK, made it an ideal reagent for the first enrichment step. To test this, we activated NS3a-CIAR in HEK293 cells transiently expressing the tdMCP-ERK2(K54R) reporter and enriched the lysate with immobilized E40, followed by enrichment of the flow-through using pE59 or pEM1. As expected, immobilized E40 captured almost all of the unphosphorylated tdMCP-ERK2(K54R) reporter with almost undetectable enrichment of the phosphorylated species (**Figure 2F** and **S1B**). Furthermore, both pE59 and pEM1 at the selected immobilization concentrations efficiently enriched tdMCP-pERK2(K54R) from the flow-through. Together, these results demonstrate that phospho-state-selective DARPins support efficient sequential enrichment of both ERK species from a single lysate, providing the basis for a ratiometric measurement of ERK activation state.

### Sequential enrichment of unphosphorylated and phosphorylated ERK species enables ratiometric pathway activity measurements

Having identified optimal DARPin concentrations for use in a sequential enrichment strategy with a signaling-inert reporter, we compared the resolution and dynamic range achievable with different pERK enrichment regimes in the LABEL-seq workflow (**Figure 3A**). To do this, we utilized a barcoded library of ∼1,600 untagged BRAF variants spanning 80 positions enriched with annotated germline and somatic variants and covering a wide range of capacities to promote pathway activation (**Figure S1C**). ^18,19^ Wild-type (WT) CRAF, which heterodimerizes with BRAF to promote downstream pathway activation, and tdMCP-ERK2(K54R) were transiently coexpressed in HEK293 cells stably expressing individual barcoded BRAF variants. Cells were lysed and lysates subjected to one of four enrichment regimes: E40 followed by a pERK antibody, E40 followed by pE59, E40 followed by pEM1, or our original enrichment using a pERK antibody normalized to total barcode frequency using an anti-FLAG enrichment. Following enrichment, captured circRNAs were reverse transcribed and quantified by NGS, and pathway activity scores were calculated for all BRAF variants.

**Figure 3:**
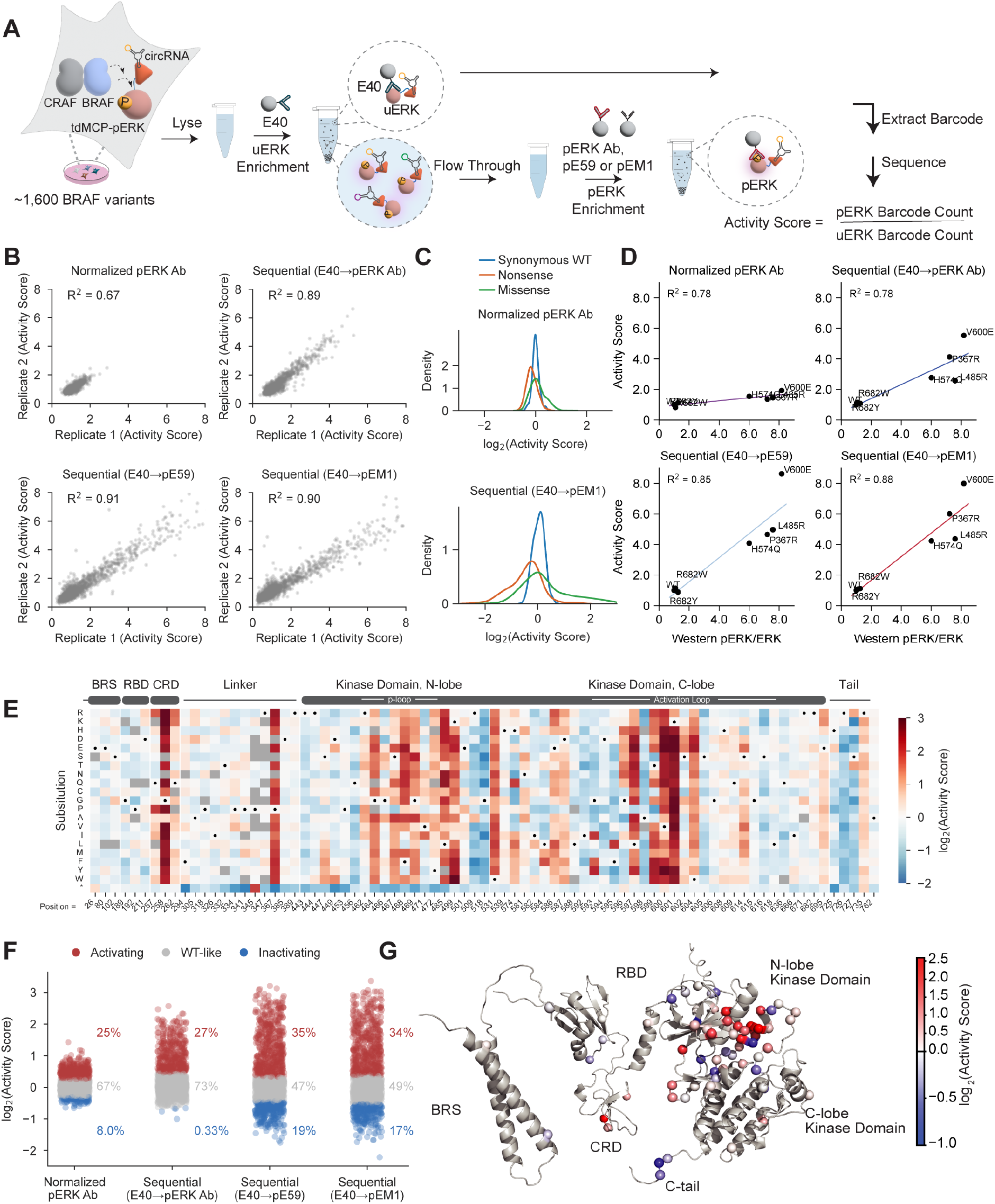
Sequential DARPin enrichment expands the dynamic range of BRAF variant activity measurements. **(A)** Schematic of the three sequential enrichment workflows used to measure BRAF variant pathway activity scores. Lysate from cells stably expressing untagged BRAF variants and transiently coexpressing WT CRAF and the tdMCP-ERK2(K54R) reporter is first depleted of uERK with immobilized E40 (4 µM); the flow-through is then enriched with one of three pERK-selective reagents: anti-pERK antibody-conjugated beads (E40 → anti-pERK antibody), immobilized pE59 (2 µM; E40 → pE59), or immobilized pEM1 (0.5 µM; E40 → pEM1). Barcodes recovered from each enrichment are sequenced, and pathway activity scores are calculated as the ratio of barcoded circRNAs captured by the pERK-selective reagent to those captured by E40. **(B)** Scatterplots comparing WT-normalized pathway activity scores from two independent enrichments performed on aliquots of a single lysate, for the normalized anti-pERK antibody enrichment and each of the three sequential enrichments in (A). Each point is one BRAF variant. R²: normalized anti-pERK antibody, 0.67; E40 → anti-pERK antibody, 0.89; E40 → pE59, 0.91; E40 → pEM1, 0.90. **(C)** Distributions of the WT-normalized log_2_ pathway activity scores for synonymous WT (blue), nonsense (orange), and missense (green) variants, obtained with normalized anti-pERK antibody enrichment and with sequential E40 → pEM1 enrichment. **(D)** Correlation of LABEL-seq and western blot measurements of pathway activation for seven individually expressed BRAF variants (WT, P367R, L485R, H574Q, R682W, R682Y, V600E). Western-derived pERK/ERK values, normalized to WT (x axis), are plotted against LABEL-seq pathway activity scores (y axis) for each of the three enrichment regimes in (A) and for the normalized anti-pERK antibody enrichment, with a regression line per regime. (n = 3 for the western quantification). **(E)** Heatmap of WT-normalized log_2_ pathway activity scores measured with sequential E40 → pEM1 enrichment in cells coexpressing WT CRAF and the tdMCP-ERK2 reporter under basal RAS-GTP. Individual amino acid substitutions (rows) are shown across the 80 profiled BRAF positions (columns); WT residues are marked with dots, and the color scale runs from −2 (inactivating) to +3 (activating). Unquantified variants are grey. (n = 2 independent E40 → pEM1 enrichments). **(F)** Functional classification of BRAF variants for each enrichment regime. Variants are colored by class (activating, red; WT-like, grey; inactivating, blue). Variants scoring more than 2σ above the synonymous WT mean were classified as activating, those more than 2σ below as inactivating, and the remainder as WT-like, with σ calculated from the matched synonymous WT distribution for each regime. Class percentages are given to the right of each column. **(G)** Position-averaged scores from (E) mapped onto an AlphaFold model of full-length BRAF (AF-P15056-F1), with the 80 profiled positions shown as spheres. Colors span a log_2_ scale of −1.0 to 2.5, with WT = 0. Annotated regions: BRAF-specific domain (BRS), RAS-binding domain (RBD), cysteine-rich domain (CRD), kinase domain, and C-terminal tail. *See also Figures S1 and S2*.

To compare the reproducibility, resolution, and dynamic range of our four enrichment strategies, we calculated replicate correlations of pathway activity scores (**Figure 3B**) and frequency distributions of the synonymous WT, missense, and nonsense variant populations (**Figure 3C**). As expected, enrichment with a pERK antibody alone provided good replicate correlation (Pearson’s R² = 0.67) but a limited range of pathway activity scores. In contrast, all three sequential enrichment regimes using uERK-selective E40 followed by a pERK-selective affinity reagent provided superior replicate correlations (Pearson’s R² ≥ 0.89 for all three; **Figure 3B**) and a broader dynamic range. Sequential DARPin enrichment of the tdMCP-ERK2 reporter using pE59 or pEM1 after E40 provided the highest resolution between the synonymous WT distribution and high- and low-activity missense variants, with scores spanning an approximately 50-fold range (**Figure S1D**). Pathway activity scores from sequential DARPin enrichments correlated well with pERK levels measured by western blot for select BRAF variants (**Figure 3D**), and fold increases for activating variants relative to WT BRAF were comparable between the two methods.

We classified a BRAF variant as activating if its pathway activity score exceeded the synonymous WT mean by more than two standard deviations, and as inactivating if it fell more than two standard deviations below this mean. Comparison of the percentage of variants classified as activating or inactivating across the four enrichment strategies (**Figure 3F**) revealed that sequential DARPin enrichments facilitated identification of a greater proportion of both activating and inactivating BRAF variants, with the largest increase observed among variants with pathway activity scores substantially below that of WT BRAF. Mapping the LABEL-seq data from the sequential enrichment strategy onto a sequence-pathway activity map (**Figure 3E** and **S2A-C**) and projecting position-averaged pathway activity scores onto an AlphaFold-generated structure of full-length BRAF (**Figure 3G**) revealed expected patterns.

For example, variants at positions known to stabilize autoinhibitory interactions, including those in the inhibitory turn of the activation loop (Leu597, Val600, and Lys601) and the phosphate-binding loop (Gly464, Phe468, and Gly469), led to high pathway activity scores. Conversely, variants at Arg509, a residue that mediates active RAF dimer formation, were almost uniformly inactivating relative to WT BRAF.

Profiling of the ∼1,600 BRAF variant library with our high-resolution pathway activity reporter allowed us to assign pathway activity scores to 70 somatic BRAF variants in The Cancer Genome Atlas (TCGA) and 39 germline variants associated with RASopathies annotated in ClinVar^20, 21^. Among TCGA variants, 55% were classified as pathway-activating and 10% as inactivating (**Figure S2D**). As expected, the Class 1 variant Val600Glu and Class 2 variant Lys601Glu^22^ possessed among the highest pathway activity scores. Somatic variants in BRAF’s phosphate-binding loop, such as Gly469Ala, Gly464Glu, and Phe468Cys, also exhibited high pathway activity scores. In contrast, most germline variants associated with RASopathies, such as Gln257Arg in the CRD and Glu501Val, displayed more modest increases in pathway activation relative to WT BRAF (**Figure S2E**). Consistent with this, only 33% of RASopathy-associated variants were classified as pathway activating. Overall, somatic TCGA variants exhibited generally higher pathway activity scores than RASopathy-associated variants in ClinVar.

### Comparison of pathway activity scores under different RAS-GTP levels reveals determinants of BRAF autoinhibition and active dimer formation

Unstimulated HEK293 cells have low levels of RAS-GTP and consequently WT BRAF is mainly in an autoinhibited state. Under these basal conditions, substitutions in BRAF that promote increased pathway activity do so primarily by disrupting autoinhibitory interactions, releasing a signaling-competent state. Because most BRAF variants were not pathway activating under basal conditions (**Figure 3F**), it is unclear whether their autoinhibition could be relieved and whether they could be recruited into active RAF dimers by RAS-GTP in the same manner as WT BRAF. We therefore measured pathway activity scores for the BRAF variant library in cells with elevated RAS-GTP levels (**Figure 4A**). To provide a comparison in which fewer dimeric RAF species need to be considered when assessing the mechanistic determinants of pathway activation, we first measured pathway activity scores for the untagged BRAF variant library in unstimulated HEK293 cells in the absence of overexpressed CRAF, including barcoded pathway activity standards to facilitate quantitative comparison with subsequent experiments (**Figure S3A, S3B**). As with coexpressed CRAF under basal conditions, pathway activity scores for the BRAF variant library alone were well correlated (**Figure S3C**) and spanned a wide range (**Figure 4B**, **C**), with activating and inactivating variants identified at similar positions (**Figure 3E**).

**Figure 4:**
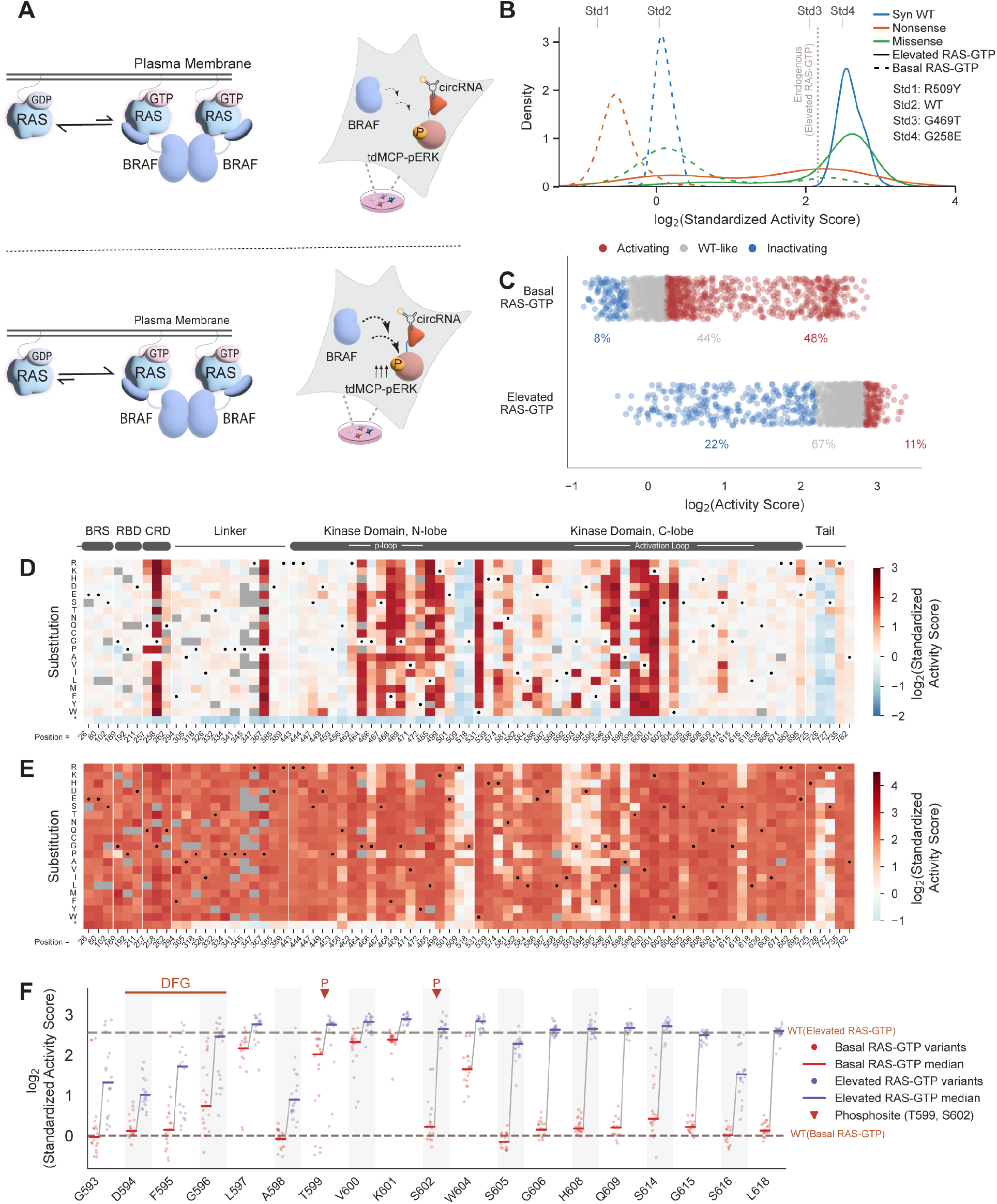
Paired profiling of BRAF variants at basal and elevated RAS-GTP levels. (**A**) Schematic of paired measurements under basal and elevated RAS-GTP levels. For the basal RAS-GTP condition (Top), the untagged BRAF variant library was integrated into the parental circRNA Protein Landing Pad (RPLP) line; for the elevated RAS-GTP condition (Bottom), the same library was integrated into an RPLP line stably expressing the NS3a-CIAR switch. Both lines were transduced with the tdMCP-ERK2 reporter. Before lysis, the NS3a-CIAR line was treated with danoprevir to raise endogenous RAS-GTP levels. Both conditions included barcoded spike-in standards, placing all scores on a common standardized scale. **(B)** Distributions of log_2_ standardized pathway activity scores for synonymous WT (blue), nonsense (orange), and missense (green) BRAF variants under elevated (solid lines) and basal (dashed lines) RAS-GTP. Vertical dashes mark the four spike-in standards and their assigned reference pathway activity scores (Std1, 0.64; Std2, 1.00; Std3, 4.05; Std4, 5.62); the vertical dotted line marks the endogenous pathway activity under elevated RAS-GTP (4.48; log_2_ = 2.16). n = 2 biological replicates per condition. **(C)** Functional classification of BRAF variants under basal and elevated RAS-GTP. Variants are colored by class (activating, red; WT-like, grey; inactivating, blue), assigned relative to WT ± 2σ of the synonymous WT distribution matched to each condition; class percentages are given at the bottom of each row. **(D)** Heatmap of log_2_ standardized pathway activity scores under basal RAS-GTP. Substitutions (rows) across the 80 profiled positions (columns); color scale −2 to +3, centered at 0; unquantified variants grey. n = 2 biological replicates. **(E)** Heatmap of log_2_ standardized pathway activity scores under elevated RAS-GTP. The scale ranges from −1 to +4 and is centered at 0. Unquantified variants are shown in grey. n = 2 biological replicates. **(F)** log_2_ standardized pathway activity scores for variants at each profiled position in BRAF’s activation loop (residues 593-623). Each dot is one variant; red, basal RAS-GTP; purple, elevated RAS-GTP. Horizontal bars give the median for each condition at each position, and lines connect medians between conditions. The DFG motif (residues 594-596) and the activation-loop phosphosites tested (Thr599, Ser602) are highlighted. *See also Figure S3*.

To assess the BRAF variant library under elevated RAS-GTP levels, we used the genetically encoded protein switch NS3a-CIAR (**Figure S1A**).^16, 17^ HEK293 cells coexpressing the untagged BRAF variant library and NS3a-CIAR were treated with danoprevir for 1 hr, lysed, and pathway activity scores were measured alongside barcoded pathway activity standards.

Replicate pathway activity scores were again well correlated (**Figure S3D**) and were on average markedly higher than those obtained under basal conditions (**Figure 4B, C**). As expected, WT BRAF promoted substantially higher pathway activation under elevated RAS-GTP levels (**Figure 4B, C**), with its standardized pathway activity score approaching that of the most activating BRAF variants under basal conditions (**Figure 4C**). Under elevated RAS-GTP levels, approximately 78% of BRAF missense variants had WT-like or greater pathway activity scores, demonstrating that most positions frequently mutated in cancer maintain the ability to form signaling-active RAF dimers. Notably, 11% of missense variants exceeded the pathway activity of WT BRAF, indicating that substitutions at some positions are able to further enhance the ability of BRAF to form active signaling complexes under elevated RAS-GTP levels.

Our BRAF variant library includes positions involved in autoinhibition and the formation of active RAF dimers, including the cysteine-rich domain (CRD), phosphate-binding and activation loops, and the regulatory C-terminal tail. Comparison of pathway activity scores at these positions under basal and elevated RAS-GTP levels provided insight into the roles these regions play in stabilizing the autoinhibited conformation and in facilitating active signaling complex formation.

Substitutions that destabilize the autoinhibited conformation increase pathway activity under basal conditions, whereas those that disrupt active signaling complex formation prevent variants from achieving WT-like or higher pathway activity under elevated RAS-GTP levels. We profiled three positions (Gln257, Gly258, and Gln262) in BRAF’s CRD, which contributes to both autoinhibition and recruitment to membrane-bound RAS-GTP (**Figure 4D, E**).^23, 24^ Most substitutions at Gly258 led to high pathway activity scores under basal conditions, whereas substitutions at the neighboring positions Gln257 and Gln262 largely exhibited WT-like pathway activity scores. In cells with elevated RAS-GTP levels, many Gly258 variants yielded pathway activity scores exceeding that of WT BRAF, whereas Gln257 and Gln262 variants promoted pathway activity to levels comparable to WT BRAF. Together, these observations suggest that Gly258 contributes to stabilization of the autoinhibited conformation but does not facilitate the formation of contacts that are required for the recruitment of BRAF to active signaling complexes at membranes, while Gln257 and Gln262 do not play essential roles in either process.

Positions in BRAF’s activation loop (residues 593-623) harbor a number of clinically observed somatic mutations, including the most frequently observed variant (Val600Glu) in cancer,^25,26^ and our BRAF variant library facilitated profiling of just over 60% of all positions in this region (**Figure 4E, F**). At five activation loop positions (Leu597, Thr599, Val600, Lys601, and Trp604), almost all substitutions resulted in markedly higher pathway activity scores than WT BRAF under basal conditions, consistent with these positions forming a network of hydrophobic interactions that stabilize the inactive conformation of BRAF’s αC helix (**Figure 4F**). In the presence of elevated RAS-GTP, almost all substitutions at these positions demonstrated WT-like or higher pathway activity scores, suggesting that these positions do not participate in interactions that are required for the formation of active signaling complex. Thr599 has been identified as a site of activation loop phosphorylation ^27^, and multiple substitutions at this position, including Asp and Glu, promoted high pathway activity under basal conditions, indicating that phosphorylation at this position would be expected to disrupt autoinhibition. However, almost all variants at Thr599 yielded pathway activity scores equal to or greater than WT BRAF in cells with elevated RAS-GTP levels, consistent with Thr599 phosphorylation not being required for BRAF to participate in active signaling dimers.

In contrast to the five activation loop positions that stabilize the autoinhibitory turn, the remaining 14 profiled positions largely lacked variants with high pathway activity scores under basal conditions. Eight of these activation loop positions (Gly596, Ser602, Gly606, His608, Gln609, Ser614, Gly615, and Leu618) also exhibited median WT-like pathway activity scores under elevated RAS-GTP levels (**Figure 4F**), suggesting that these positions do not participate in autoinhibition or the formation of active BRAF signaling complexes. One of these positions, Ser602, has also been identified as a site of activation loop phosphorylation. ^28^ In contrast to Thr599, only the phosphomimetic substitutions Asp and Glu at Ser602 led to high pathway activity scores in the presence of basal RAS-GTP, supporting the notion that it is the negative charge introduced by phosphorylation, rather than alteration of the WT residue, that leads to disruption of autoinhibition at this site. Similar to Thr599, most Ser602 variants had WT-like pathway activity scores under elevated RAS-GTP levels, consistent with Ser602 phosphorylation not being required to form active RAF dimers. The remaining six positions in BRAF’s activation loop had median pathway activity scores below that of WT BRAF in cells with elevated RAS-GTP levels, suggesting that the WT residue at these positions contributes to active BRAF dimer formation.

### Many BRAF variants exert dominant negative effects on pathway activity

The high resolution and broad dynamic range of the tdMCP-ERK2 reporter allowed us to identify BRAF variants with pathway activity scores substantially lower than WT BRAF under basal conditions (**Figures 4D** and **5A**), indicative of their exertion of a dominant negative effect on pathway activation. Analysis of standardized pathway activity scores in cells with elevated RAS-GTP levels revealed that many of these same variants suppressed pathway activation well below endogenous levels (**Figures 4B** and **5A**), with some variants causing dramatic reductions indicative of strong dominant negative activity. To validate these findings, we selected representative dominant negative BRAF variants and assessed their effect on endogenous ERK activation in cells with elevated RAS-GTP levels by western blot analysis (**Figures 5B** and **5C**). Exogenous expression of all selected variants led to markedly lower levels of pERK than cells expressing WT BRAF (**Figure 5B**) or an empty vector (**Figure 5C** and **S4A**), and the magnitude of this reduction correlated with pathway activity scores (**Figure S4B**). Several variants (R509W, G518T, S727K, and R726D) reduced pERK to almost undetectable levels despite the high pathway activation in cells with elevated RAS-GTP levels. Having confirmed the dominant negative effects of individual variants, we systematically assessed the prevalence of this phenomenon across all profiled positions by calculating position-averaged pathway activity scores in cells with elevated RAS-GTP. Of the 80 profiled positions, 16 had average pathway activity scores below endogenous levels in cells with elevated RAS-GTP levels, demonstrating that the WT residue at a substantial number of positions within BRAF actively suppresses an intrinsic tendency toward dominant negative behavior that is unmasked upon substitution (**Figure S4C)**. These positions are distributed throughout BRAF’s kinase domain, including within and surrounding the activation and phosphate-binding loops, αC-helix, and C-terminal tail (**Figure 5D**).

**Figure 5:**
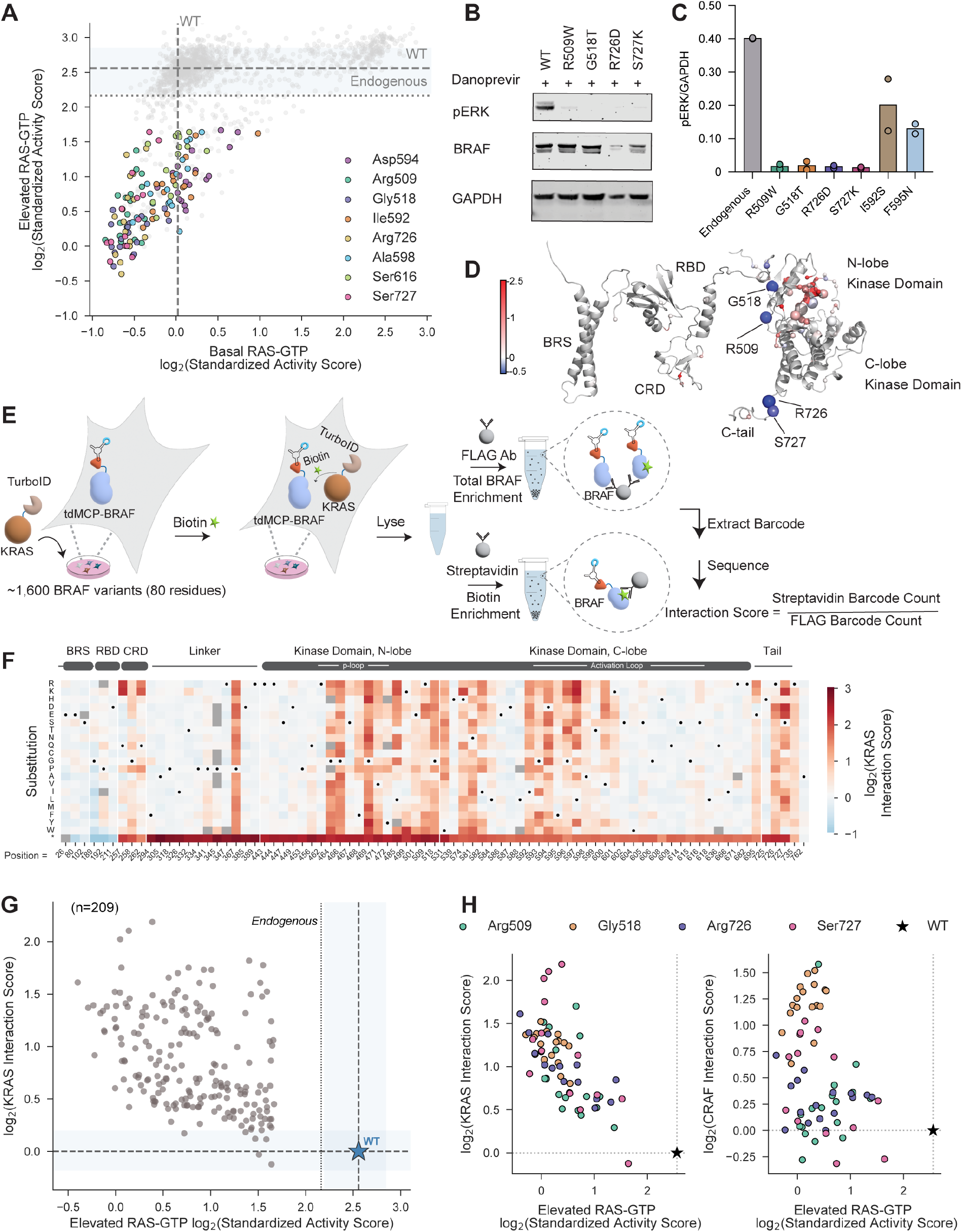
Identification of dominant negative BRAF variants and their engagement of KRAS. **(A)** log_2_ standardized pathway activity scores under elevated RAS-GTP plotted against scores under basal RAS-GTP. Vertical dashed line, WT activity under basal RAS-GTP; horizontal dashed line, WT activity under elevated RAS-GTP; horizontal dotted line, endogenous pathway activity under elevated RAS-GTP in the absence of exogenous BRAF. Grey dots, individual variants; colored dots, variants at the eight positions most enriched for dominant negative substitutions (Arg509, Gly518, Ile592, Asp594, Ala598, Ser616, Arg726, Ser727). **(B)** Representative western blot of endogenous pERK in HEK293 cells stably expressing NS3a-CIAR together with WT BRAF or one of the dominant negative variants shown. Cells were treated with danoprevir to raise endogenous RAS-GTP levels prior to lysis (n = 2 biological replicates). **(C)** Quantification of GAPDH-normalized endogenous pERK in HEK293 lines stably expressing NS3a-CIAR together with either the dominant negative variants shown or an empty vector, all treated with danoprevir to raise endogenous RAS-GTP levels; blot shown in Figure S4A (n = 2 biological replicates). **(D)** Position-averaged standardized pathway activity scores under basal RAS-GTP mapped onto the AlphaFold model of BRAF (AF-P15056-F1), with the 80 profiled positions as spheres. Colors span a log_2_ scale of −0.5 to 2.5 centered at WT activity. Sphere radius scales linearly with the percentage of dominant negative variants at each position. Annotated regions as in Figure 3G. **(E)** Workflow of the LABEL-seq interaction assay. Cells coexpressing 3xFLAG-tdMCP-BRAF variants and transduced with TurboID-KRAS are labeled with biotin; the lysate is split for anti-FLAG and streptavidin enrichment, followed by barcode sequencing. Interaction scores are the ratio of streptavidin-to FLAG-enriched barcodes, normalized to WT within each replicate. **(F)** Heatmap of WT-normalized log_2_ KRAS interaction scores across the BRAF variant library, measured as described in (E). Substitutions (rows) across profiled positions (columns); WT residues marked with dots; color scale −1 to +3, centered at WT (log_2_ = 0); unquantified variants are grey. n = 2 biological replicates. **(G)** KRAS interaction score (log_2_) versus elevated RAS-GTP standardized pathway activity score (log_2_) for the 209 variants classified as dominant negative. Shaded region, WT range; star, WT control. **(H)** WT-normalized KRAS (left) and CRAF (right) interaction scores (log_2_) versus elevated RAS-GTP standardized pathway activity scores (log_2_) for dominant negative variants at Arg509, Gly518, Arg726, and Ser727. Star represents WT. CRAF interaction scores are adapted from Simon et al. ^5^ *See also Figure S4*.

The observation that almost all truncating nonsense BRAF variants possessing a complete RBD-CRD module but lacking an intact kinase domain were dominant negative (**Figure 4E**) led us to hypothesize that many dominant negative missense variants share similar properties: enhanced recruitment to membrane-bound RAS relative to WT full-length BRAF, where they potentially outcompete endogenous RAF kinases, coupled with an inability to form active signaling complexes.^29,30^ To test this, we measured the interaction of the BRAF variant library with WT KRAS using a LABEL-seq proximity labeling assay in which a tdMCP-tagged BRAF variant library was coexpressed with TurboID-KRAS (**Figure 5E**)^31^, allowing interactions to be captured by proximity-dependent biotinylation. Interaction scores for each variant relative to WT BRAF were calculated following biotin labeling and streptavidin enrichment. Replicate interaction scores were well correlated (**Figure S4D**) and spanned a wide range of BRAF variant-KRAS interaction levels (**Figure 5F**). As expected, BRAF nonsense variants with a complete RBD-CRD module but lacking an intact kinase domain showed substantially higher KRAS interaction scores than full-length WT BRAF (**Figure 5F**). Supporting our hypothesis, most dominant negative missense variants displayed increased KRAS interaction relative to WT BRAF, with stronger dominant negative effects trending with higher KRAS interaction scores (**Figure 5G**). Together, these findings suggest a mechanism by which many BRAF missense variants exert dominant negative effects: disruption of autoinhibitory interactions that normally prevent recruitment to membrane-bound RAS, coupled with an inability to promote downstream pathway activation following recruitment.

The four positions with variants that exerted the largest dominant negative effects under both basal and elevated RAS-GTP levels were Arg509, Gly518, Arg726, and Ser727 (**Figure 5A**). Arg509 is located directly C-terminal to BRAF’s αC-helix, where it forms part of the interprotomer interface of RAF dimers, and substitutions at this position have been shown to weaken RAF dimerization. ^5,32,33^ Our interaction profiling revealed that substitutions at Arg509 also promoted increased interactions with KRAS (**Figure 5H**), most likely by disrupting autoinhibitory interactions through an as yet unknown mechanism. Together, these observations support a model in which variants at Arg509 exert their dominant negative effect by outcompeting endogenous RAF kinases for recruitment to active RAS while retaining a reduced ability to form RAF homodimers or heterodimers. Gly518 directly precedes the β4-strand in BRAF’s N-lobe and is located in close proximity to the interprotomer interface of RAF dimers. Like Arg509, most Gly518 substitutions promoted increased interactions with KRAS, suggesting that they similarly outcompete endogenous RAF kinases for RAS binding. However, unlike Arg509, substitutions at Gly518 were previously found to lead to increased interactions with CRAF (**Figure 5H and S4E**)^5^, indicating dominant negative variants at this position are capable of forming RAF dimers upon recruitment to active RAS but that neither protomer of the resulting complex is catalytically active.

Arg726 and Ser727 are part of BRAF’s C-terminal tail, which contains a constitutively phosphorylated residue (Ser729) that promotes high-affinity interaction with one protomer of 14-3-3 dimers. Both the kinase believed to phosphorylate Ser729 (AMPK) and 14-3-3 proteins show a strong preference for Arg and Ser at the −3 and −2 positions relative to Ser729 ^34,35^ suggesting that dominant negative variants at Arg726 and Ser727 likely lack both C-terminal tail phosphorylation and phospho-Ser729-mediated 14-3-3 interactions. Consistent with previous studies demonstrating that interaction of BRAF’s C-terminal tail with 14-3-3 dimers facilitates both autoinhibition and active RAF dimer formation ^36,37, 38, 39, 40, 41^, dominant negative variants at Arg726 and Ser727 demonstrated increased KRAS interactions and decreased CRAF interactions (**Figure 5H**). This suggests that loss of 14-3-3 binding simultaneously disrupts the autoinhibitory mechanisms that limit RAS engagement and impairs productive RAF dimer formation, thereby producing a dominant negative effect (**Figure 5A**). Together, these findings reveal that dominant negative behavior in BRAF arises from a common mechanistic theme involving disruption of autoinhibitory constraints that normally limit recruitment to RAS, combined with an impaired capacity to form or adopt catalytically active RAF dimers, and that this susceptibility is encoded at structurally distinct positions distributed throughout the kinase domain.

### Profiling drug responses with LABEL-seq reveals variant-specific sensitivity and resistance to the ATP-competitive inhibitor LXH254

We next used our LABEL-seq tdMCP-ERK2 reporter assay to measure the susceptibility of BRAF variants to inhibition by LXH254 (naporafenib), a highly selective ATP-competitive pan-RAF inhibitor that stabilizes the DFG-out conformation^42^ and is currently undergoing clinical evaluation for RAS/MAPK-altered solid tumors and melanoma.^43,44^ We reasoned that our assay could be used to measure the intracellular potency of LXH254 in blocking pathway activation promoted by multiple clinically relevant BRAF variants and to identify sequence determinants of drug sensitivity and resistance. To do this, we treated cells expressing the BRAF variant library and the tdMCP-ERK2 reporter with a dose range of LXH254 concentrations and measured the pathway activity score of each variant at each concentration (**Figure 6A, 6B**, **S5A**, and **S5B**).

**Figure 6:**
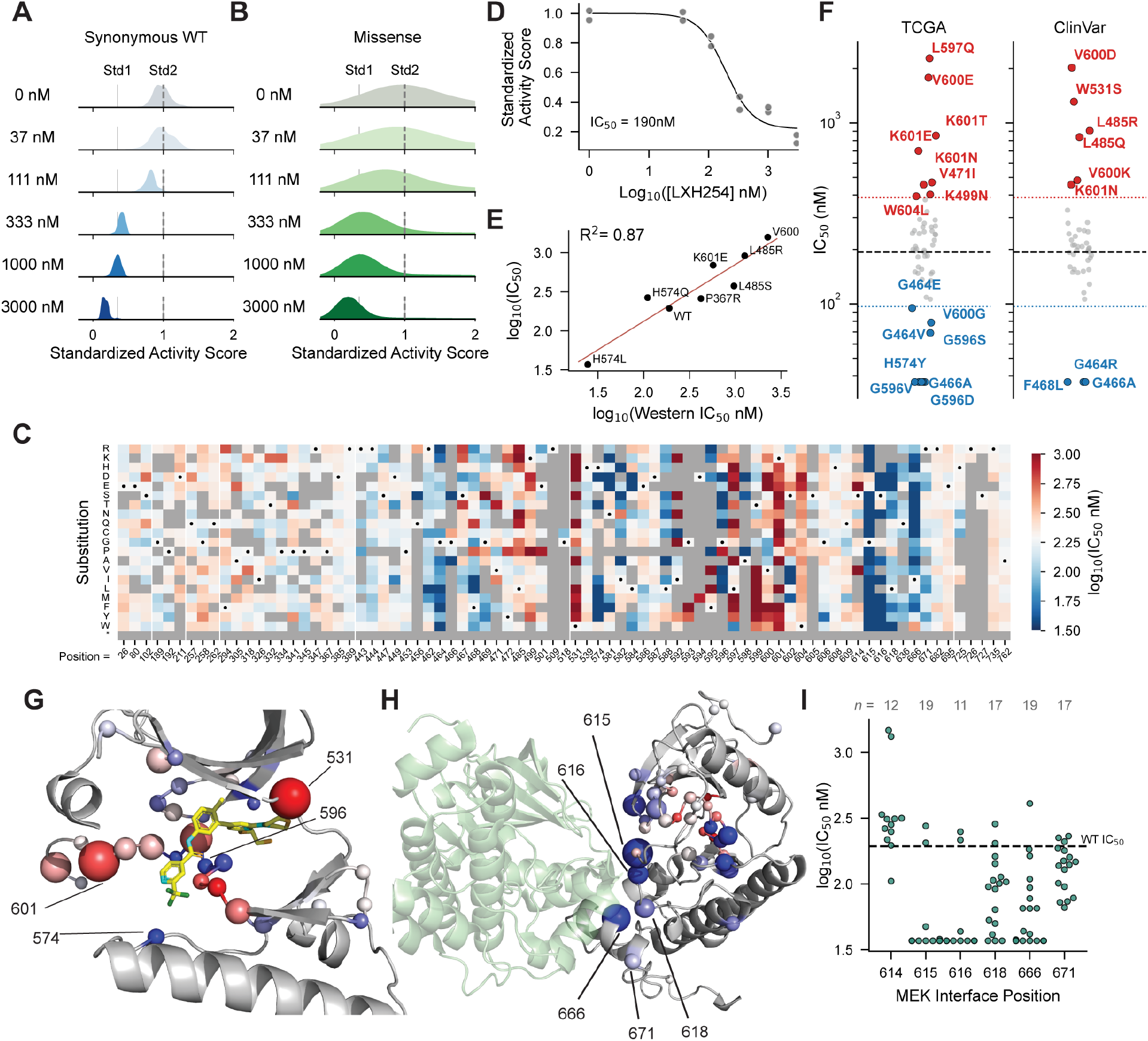
**Intracellular LXH254 potency and its structural determinants across ∼1,000 BRAF variants**. **(A)** Distributions of standardized activity scores for synonymous WT variants after 4 h of treatment with LXH254 at five concentrations (37, 111, 333, 1,000, and 3,000 nM) or with DMSO. Vertical lines mark normalization standards. n = 2 biological replicates per condition. **(B)** As in (A), for missense variants. **(C)** Heatmap of IC_50_ values across the 80 profiled positions and the 19 amino acid substitutions at each position. Color encodes log_10_(IC_50_, nM) on a red-blue diverging scale (red, resistant; blue, sensitizing), centered on the WT IC_50_ (190 nM; log_10_ = 2.28). Black dots mark WT residues. Fits falling below or above the tested concentration range were assigned IC_50_ values of 37 and 3,000 nM, respectively; variants for which an IC_50_ value could not be assigned are shown in grey. **(D)** Representative dose-response curve for WT BRAF (IC_50_ = 190 nM). **(E)** Correlation (R² = 0.87) of IC_50_ values derived from standardized pathway activity scores with IC_50_ values derived from western blot quantification, for the individually expressed BRAF variants shown (blots in Figure S5C). **(F)** IC_50_ values for BRAF variants annotated in TCGA and in ClinVar, grouped by database of origin and colored by classification (resistant, red; WT-like, grey; sensitizing, blue). Dashed black line, WT IC_50_; dashed red line, 2-fold resistance threshold; dashed blue line, 2-fold sensitization threshold. **(G)** Positional-median LXH254 IC_50_ values for W531, H574, G596, and K601, mapped onto the crystal structure of the BRAF dimer bound to LXH254 (PDB: 8F7P). Color denotes median IC_50_ (red, resistant; blue, sensitizing); sphere radius scales with the number of resistant variants at each position. **(H)** Positional-median LXH254 IC_50_ values mapped onto a cryo-EM structure of the autoinhibited BRAF-MEK1 complex (PDB: 6NYB), with MEK1 in forest green and BRAF in grey. Sphere size indicates the number of sensitizing variants at each position; color denotes median IC_50_ value at each position. Labeled BRAF positions are within 4.5 Å of MEK1 in the autoinhibited complex. **(I)** IC_50_ values for individual substitutions at the six profiled BRAF-MEK1 contact positions. Each point is one substitution; dashed line, WT IC_50_. *See also Figure S5*.

Barcoded pathway activity standards were included with all conditions to enable quantitative comparison of standardized pathway activity scores across LXH254 concentrations (**Figure S3A**). Synonymous WT BRAF variants showed a dose-dependent decrease in pathway activity scores (**Figure 6A**), with no paradoxical activation observed at any concentration tested, consistent with the reported ability of LXH254 to promote RAF dimerization without negative cooperativity between protomers.^45, 46^ Missense variants also displayed dose-dependent decreases in pathway activity, but the magnitude of inhibition and the concentration at which it occurred varied widely across variants (**Figure 6B**).

The standardized pathway activity scores at each LXH254 concentration were used to calculate half-maximal inhibitory concentration (IC_50_) values for each variant (**Figure 6C, 6D**, and **S5B**). We were able to assign IC_50_ values to ∼ 1,000 BRAF variants, ranging from less than 37 nM, the lowest IC_50_ value that could be estimated based on the LXH254 concentrations tested, to greater than 3,000 nM, with WT BRAF possessing an intracellular IC_50_ value of 190 nM (**Figure 6D**). IC_50_ values calculated from pathway activity scores correlated well (R^2^ = 0.87) with those determined using western blot analysis for select BRAF variants (**Figure 6E** and **S5C**). A majority of BRAF variants for which IC_50_ values could be determined (74%) demonstrated sensitivities to LXH254 within two-fold of WT BRAF (IC_50_ = 95-380 nM).

However, 11% of BRAF variants were more than two-fold more sensitive than WT BRAF, and 15% were more than two-fold more resistant. Among the BRAF variants annotated in TCGA and as RASopathy-associated in ClinVar, several showed increased sensitivity to LXH254 relative to WT BRAF, including variants in the phosphate-binding loop (Gly464, Gly466, and Phe468) and the activation loop (Gly596 and Val600) (**Figure 6F**).

Several positions in BRAF appeared particularly prone to resistance, with a majority of variants at these positions demonstrating reduced sensitivity to LXH254 relative to WT BRAF (**Figure 6G**). Trp531 and Lys601 had both the largest number of LXH254-resistant variants and the highest position-averaged IC_50_values among all profiled positions. Trp531 is located within the hinge region connecting the N- and C-lobes of BRAF’s kinase domain, and its indole side chain, which sits at the top of the ATP-binding site, participates in hydrophobic interactions with the morpholine moiety of LXH254. Almost all substitutions at Trp531 promoted high pathway activity under basal conditions (**Figure 4D** and **S5A**), indicating that Trp531 contributes to autoinhibition. The high resistance of W531 variants likely reflects a combination of less favorable hydrophobic interactions with LXH254 and the higher drug concentrations required to suppress their elevated basal pathway activation. Resistance to the ATP-competitive tyrosine kinase inhibitor dasatinib was previously observed at the equivalent position (Tyr343) in SRC kinase, demonstrating that variation at this hinge position represents a conserved route to kinase inhibitor resistance.^47^ Lys601 is located within BRAF’s activation loop, five positions C-terminal to the DFG motif, and unlike Trp531, its side chain does not directly contact LXH254 (**Figure 6G**). The high resistance of Lys601 variants therefore likely reflects a combination of their elevated basal pathway activation and a negative influence on the ability of BRAF’s activation loop to adopt the DFG-out conformation required for LXH254 binding. Interestingly, variants at the neighboring position Val600, which also does not contact LXH254, promoted similarly high basal pathway activation but showed more variable sensitivities to LXH254, with several demonstrating substantially higher sensitivity than WT BRAF (**Figure S5D**). Thus, the uniformly negative effect of Lys601 substitutions on the ability of BRAF’s activation loop to accommodate LXH254 is not shared by substitutions at Val600, indicating that variants in this region can exert variable conformational effects.

Like Trp531 and Lys601, His574 and Gly596 line the ATP-binding site of BRAF (**Figure 6G**), and variants at these positions promoted high pathway activity under basal conditions (**Figure 4D**). However, a majority of variants at these two positions for which IC_50_ values could be determined were substantially more sensitive to LXH254 than WT BRAF. His574 lies within BRAF’s catalytic loop, and its imidazole side chain forms interactions with Phe595 of the DFG motif in the autoinhibited BRAF complex (**Figure S5E**); Phe595 must rotate to the “out” position to accommodate the 2-trifluoromethylpyridine group of LXH254. In the DFG-out conformation, His574 sits directly below this group (**Figure S5F**). The increased sensitivity of His574 variants may therefore arise from a combination of destabilization of the interactions that maintain the DFG-in conformation and the potential of substituted residues to form more favorable hydrophobic interactions with LXH254, particularly those with large aliphatic or aromatic side chains. Position 596 is the Gly of the DFG motif within the activation loop, and substitutions at this position are unlikely to contact LXH254 directly (**Figure S5F**). Substitutions at Gly596 may therefore sensitize BRAF to LXH254 exclusively through destabilization of the DFG-in conformation. Together, these results demonstrate that substitutions at positions within or adjacent to the activation loop can confer a wide range of sensitivities to LXH254, likely reflecting differences in the relative stabilities of the DFG-in and DFG-out conformations.

A general mechanism of sensitizing BRAF to LXH254 appears to be disruption of BRAF-MEK1 interactions (**Figures 6H**, **6I**). We profiled six of the 25 positions in BRAF that form its interface with MEK1 in the autoinhibited complex and observed that sensitizing variants were highly enriched at these positions. Five of the six (Gly615, Ser616, Leu618, Ile666, and Arg671) possessed multiple LXH254-sensitized variants. Substitutions at Gly615 and Ser616 were particularly sensitizing, with almost all variants at these positions displaying IC_50_ values substantially lower than WT BRAF. Gly615 and Ser616 are part of a short segment in BRAF’s activation loop that forms an antiparallel β-strand interaction with MEK1’s activation loop, ^40^ stabilizing the αC-helix-out conformation of both kinases in the autoinhibited complex, in which BRAF’s Phe595 adopts the “in” conformation. Substitutions at Gly615 and Ser616 likely destabilize this interaction, facilitating adoption of the DFG-out conformation required for LXH254 binding ^48^. These results show that destabilizing BRAF-MEK interactions is generally sensitizing and highlight how disrupting protein-protein interactions that stabilize a kinase conformation incompatible with inhibitor binding can enhance drug sensitivity.

## Discussion

Here, we developed a high-resolution, sequencing-based assay for quantifying ERK activation state and used it to systematically profile how BRAF variation affects MAPK pathway activity. This assay’s resolution was enabled by replacing antibody-based capture of the reporter with a sequential enrichment strategy employing phospho-state-selective DARPins: the more abundant unphosphorylated tdMCP-ERK2 reporter is depleted first, and the phosphorylated reporter is then captured from the flow-through. This ratiometric approach, combined with a catalytically inert reporter that minimally perturbs cell signaling, expanded the dynamic range of pathway activity measurements and improved reproducibility relative to antibody enrichment alone. The most consequential improvement was in the identification of variants with pathway activity below that of WT BRAF, a regime that is compressed and poorly resolved by antibody-based capture. The ability to resolve low levels of pathway activation enabled the discovery of a large class of BRAF variants that suppress pathway activity below endogenous levels and that would have been misclassified as simple loss-of-function variants by lower-resolution measurements. This resolution also allowed us to profile the potency of the ATP-competitive, pan-RAF inhibitor LXH254 (naporafenib) against nearly 1,000 BRAF variants in parallel, a measurement complicated by the low activity of many of these variants under basal conditions.

Our efforts illustrate that a limiting factor often encountered in obtaining high-resolution signaling measurements is the quality of the reagents used to read out pathway state. For pathway activity assays using the LABEL-seq platform, the affinity and selectivity of the reagent used to capture a post-translationally modified form of a reporter protein that serves as a proxy for pathway activity are the major determinants of the resolution that can be obtained. While high-quality antibodies are available for enriching some post-translationally modified proteins, engineered binding proteins with defined, tunable affinities are highly attractive alternatives, ^12,49^ and the advent of new computational methods for identifying such binders will likely accelerate the development of these reagents. Here, we used affinity reagents identified through selection-based methods that recognize both the unmodified and modified forms of a reporter for sequential pulldowns; going forward, computational approaches for generating binders selective for the unmodified state could further expand this strategy.^50, 51,52^ The availability of such paired reagents will increase the breadth and diversity of signaling pathways that can be probed with high resolution using LABEL-seq.

A feature of our study was the measurement of the pathway activities of the same variants in two defined signaling contexts. Using a chemical genetic switch that facilitates an increase in endogenous RAS-GTP levels with a functionally inert small molecule, we profiled ∼1,600 BRAF variants in cells with low basal and elevated RAS-GTP levels, with barcoded pathway activity standards enabling quantitative comparison between the two conditions. This paired comparison allowed us to identify BRAF variants with disrupted autoinhibition based on their high pathway activity scores under basal RAS-GTP levels, and to determine which WT residues are required for the formation of fully functional signaling complexes at membranes under elevated RAS-GTP levels. That such assignments can be made systematically across an entire variant library highlights how pairing chemical genetic control of pathway state with precise, multiplexed measurement of protein variants is a promising strategy for dissecting how individual positions and regions within a protein contribute to multiple steps in cell signaling.

Together, our results show that high-resolution, sequencing-based readouts of pathway state can resolve functional distinctions that lower-resolution measurements obscure, separating variants that suppress signaling below endogenous levels from those that are simply inactive, and assigning intracellular inhibitor potencies across an entire variant library in a single experiment.^53^ Going forward, we envision extending this approach to other signaling pathways as paired affinity reagents selective for the modified and unmodified states of pathway-specific reporters become available. Combined with chemical genetic control of signaling state and larger variant libraries, such measurements should yield quantitative, mechanistic maps of how sequence variation shapes both signaling output and drug response across the pathways most frequently altered in disease.^54^

## Limitations of the study

Applying our MAPK pathway reporter to a BRAF variant library provided insight into the mechanisms of BRAF regulation and drug sensitivity, but our study has several limitations. All pathway activity measurements were made in cells that retain endogenous WT BRAF and CRAF, allowing the formation of multiple heterodimeric complexes between variant and WT protomers and complicating mechanistic interpretation of some pathway activity scores. In addition, all BRAF variants were exogenously overexpressed, and it remains unclear whether certain effects, such as dominant negative suppression of pathway activity, would be observed at endogenous expression levels. Finally, although our reporter is compatible with multiplexed variant libraries, it does not provide the temporal resolution of live-cell imaging approaches such as FRET reporters. Some temporal information can be obtained by lysing cells at a series of time points before reporter enrichment, but these measurements are not equivalent to continuous, real-time readouts in single cells.

## Author contributions

A.F.A., J.J.S., D.M.F., and D.J.M. conceived the study. A.F.A. developed the methodology, purified the DARPins, analyzed the data, and generated the visualizations. A.F.A. and J.J.S. planned and performed the experiments and generated and validated the cell lines. F.B.-M. provided experimental support. A.F.A. and D.J.M. wrote the original manuscript draft. All authors reviewed and edited the manuscript.

## Acknowledgments

We thank M. Walsh (UW, Seattle) and A. Zhang (UW, Seattle) for feedback on the manuscript and S. Pendyala for demuxing sequencing data. Research reported in this publication was generated using the Department of Laboratory Medicine and Pathology Flow Cytometry Core (CC101467) at the University of Washington. The results shown here are in whole or part based upon data generated by the TCGA Research Network. This work was supported by the National Institutes of Health (RM1HG010461, UM1HG011969, and R35GM152106 to D.M.F.; R01GM145011 and R01GM086858 to D.J.M.). The content is solely the responsibility of the authors and does not necessarily represent the official views of the National Institutes of Health.

---

**Figure S1.**
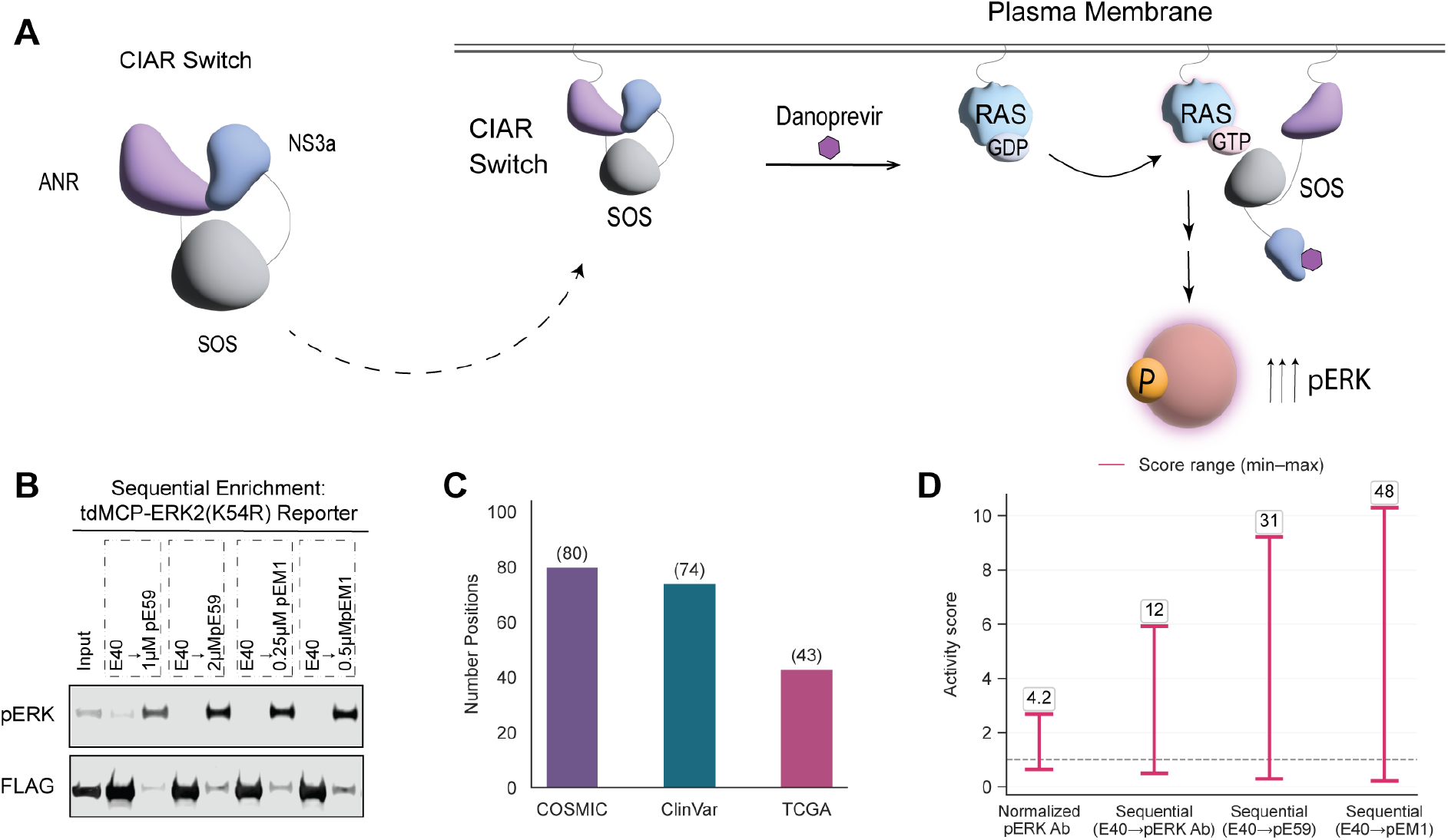
Reporter validation and BRAF library composition, related to Figures 2 and 3. **(A)** Schematic of the NS3a-CIAR system used to increase cellular RAS-GTP. The membrane-localized NS3a-CIAR switch (ANR-SOS-NS3a) contains the catalytic domain of SOS1 in an autoinhibited configuration. Danoprevir treatment relieves autoinhibition, enabling conversion of RAS-GDP to RAS-GTP. The resulting increase in endogenous RAS-GTP activates downstream MAPK signaling and increases ERK phosphorylation. **(B)** Western blot showing sequential depletion of unphosphorylated ERK (uERK) followed by enrichment of phosphorylated ERK (pERK) from lysates of NS3a-CIAR cells transiently expressing FLAG-tdMCP-ERK2(K54R) reporter. Cells were treated with danoprevir (1 µM, 1 hr.) before lysis to activate NS3a-CIAR, elevate RAS-GTP, and increase reporter phosphorylation. Lysates were first incubated with immobilized E40 DARPin (4 µM) to deplete uERK. pERK was subsequently enriched from the E40 flow-through using immobilized pE59 DARPin (1 or 2 µM) or pEM1 DARPin (0.25 or 0.5 µM), as indicated. (n = 2 biological replicates) See also Figure 2F. **(C)** Coverage of the 80 profiled BRAF positions in clinical variant databases. At least one variant was annotated at all 80 positions in COSMIC (100%), at 74 positions in ClinVar (92.5%), and at 43 positions in TCGA (53.8%). Databases were accessed as described in the STAR Methods. **(D)** Minimum-to-maximum ranges of pathway-activity scores across ∼1,600 BRAF variants for each enrichment regime in Figure 3A, and normalized pERK enrichment. Endpoints denote the lowest and highest measured scores for each assay. Labels above each range indicate the fold dynamic range, defined as the score of the highest-scoring variant divided by that of the lowest-scoring variant within each regime.

**Figure S2.**
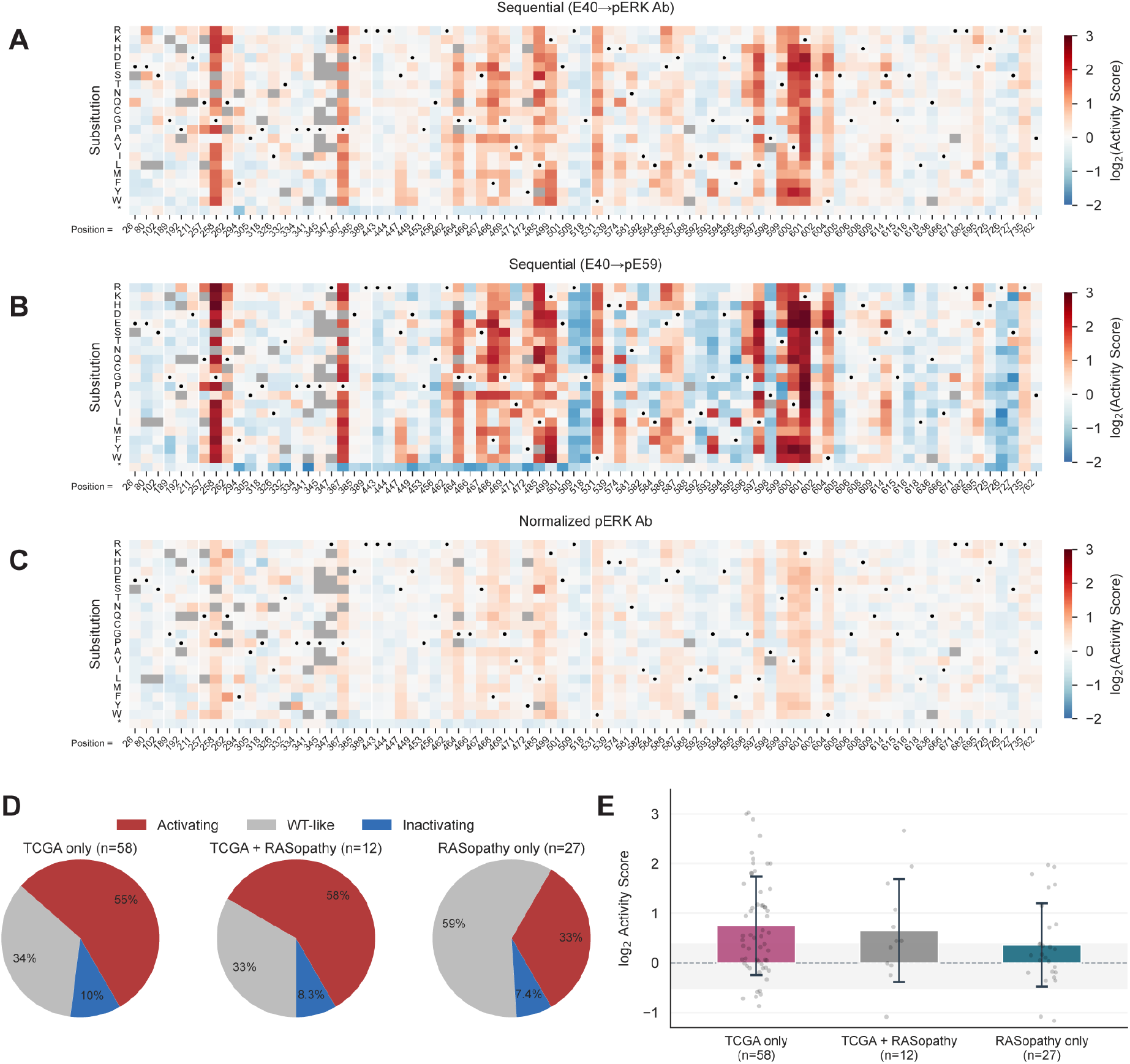
Alternative enrichment regimes and clinically annotated variants, related to Figure 3. **(A)** Heatmap of WT-normalized log_2_ pathway activity scores measured with sequential E40 → pERK Ab enrichment in cells coexpressing WT CRAF and the tdMCP-ERK2 reporter under basal RAS-GTP. Individual amino acid substitutions (rows) are shown across the 80 profiled BRAF positions (columns); WT residues are marked with dots, and the color scale runs from −2 (inactivating) to +3 (activating). Unquantified variants are grey. (n = 2 independent E40 → pERK Ab enrichments). **(B)** Heatmap of WT-normalized log_2_ pathway activity scores measured with sequential E40 → pE59 enrichment in cells coexpressing WT CRAF and the tdMCP-ERK2 reporter under basal RAS-GTP. Individual amino acid substitutions (rows) are shown across the 80 profiled BRAF positions (columns); WT residues are marked with dots, and the color scale runs from −2 (inactivating) to +3 (activating). Unquantified variants are grey. (n = 2 independent E40 → pE59 enrichments). **(C)** Heatmap of WT-normalized log_2_ pathway activity scores measured with normalized pERK enrichment in cells coexpressing WT CRAF and the tdMCP-ERK2 reporter under basal RAS-GTP. Individual amino acid substitutions (rows) are shown across the 80 profiled BRAF positions (columns); WT residues are marked with dots, and the color scale runs from −2 (inactivating) to +3 (activating). Unquantified variants are grey. (n = 2 independent normalized pERK enrichments). **(D)** Functional classification of BRAF variants annotated in TCGA and/or associated with RASopathies. Variants were classified as activating (red), WT-like (grey), or inactivating (blue) based on pathway-activity scores relative to matched synonymous wild-type variants for each enrichment regime. Scores more than 2σ above or below the synonymous wild-type mean were classified as activating or inactivating, respectively; all remaining variants were classified as WT-like. Pie charts indicate the percentage of variants in each annotation group assigned to each functional class. **(E)** Distribution of activity scores across clinical classes in (D). Bar plots show the mean log_2_ sequential E40 → pEM1 activity score for variants classified as TCGA only, TCGA + RASopathy, or RASopathy only. Each dot represents an individual variant, and error bars indicate standard deviation.

**Figure S3.**
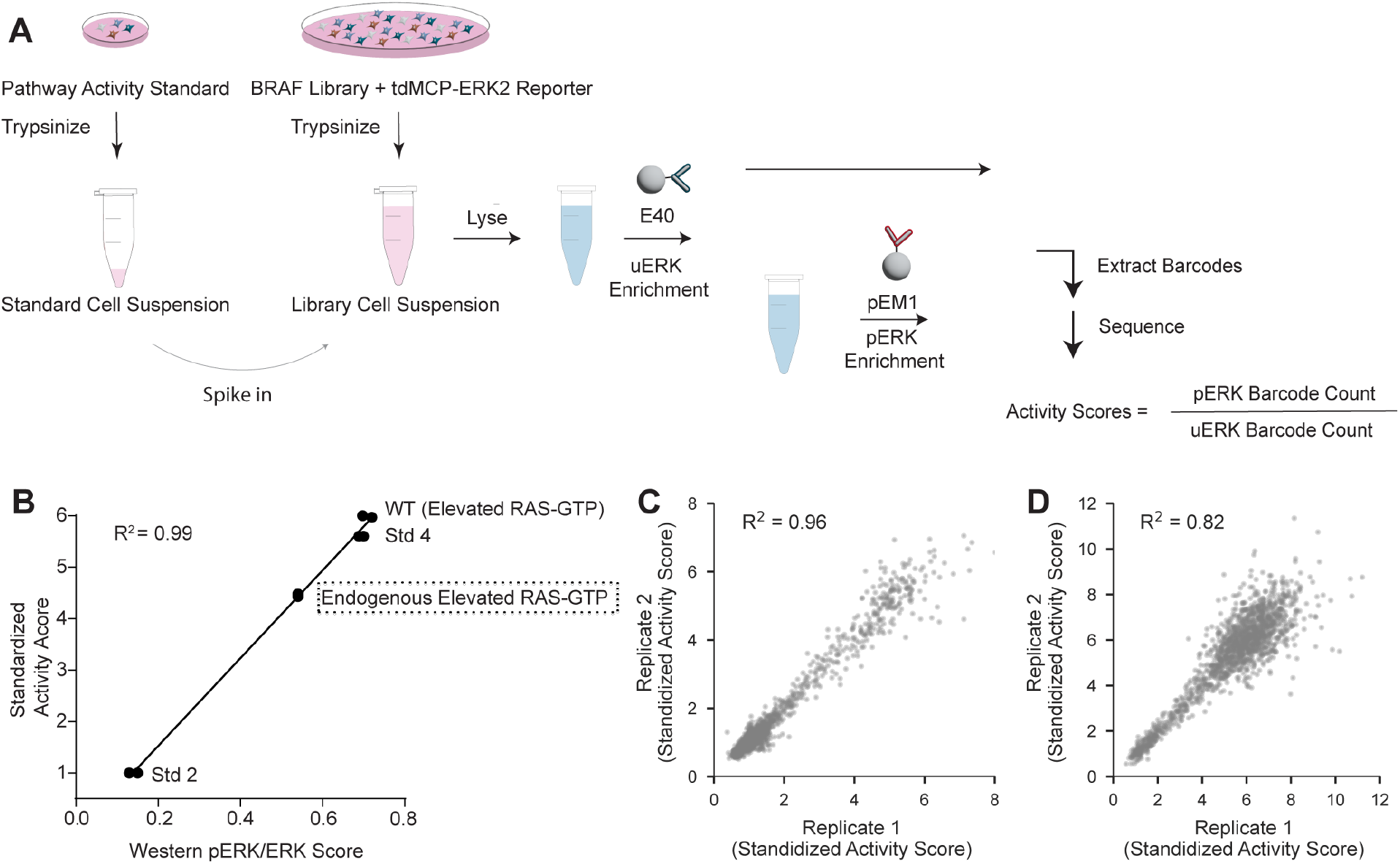
Activity standards and replicate reproducibility, related to Figure 4. **(A)** Sequential-enrichment workflow with internal standards. Orthogonally barcoded standard cell populations with distinct known pERK levels were cultured, trypsinized, and resuspended separately before being spiked into the BRAF variant-library cell suspension immediately before lysis. Mixed lysates underwent sequential uERK depletion with E40 DARPin and pERK enrichment from the flow-through with pEM1 DARPin. Barcode sequencing of the enriched fractions quantified variant and standard enrichment. Assigned standard scores were used to calibrate activity scores and assign standardized activity scores to BRAF variants. **(B)** Endogenous pathway activation under elevated RAS-GTP. HEK293 cells expressing spike-in standards (Std2 or Std4) were maintained under basal RAS-GTP conditions, whereas clonal NS3a-CIAR cell lines expressing WT BRAF or empty vector had elevated RAS-GTP. All cells were transfected with the kinase-dead ERK reporter and analyzed by western blot. Standardized activity score correlated strongly with pERK/total ERK ratios for Std2, Std4, and WT BRAF (R² = 0.99). The activity score in NS3a-CIAR empty-vector cells reflects endogenous pathway activation without exogenous BRAF. n = 2 biological replicates. **(C)** Scatterplots comparing standardized pathway activity scores from biological replicates performed under basal Ras-GTP conditions (Pearson’s R² = 0.96). **(D)** Scatterplots comparing standardized pathway activity scores from biological replicates performed under elevated Ras-GTP conditions (Pearson’s R² = 0.82).

**Figure S4.**
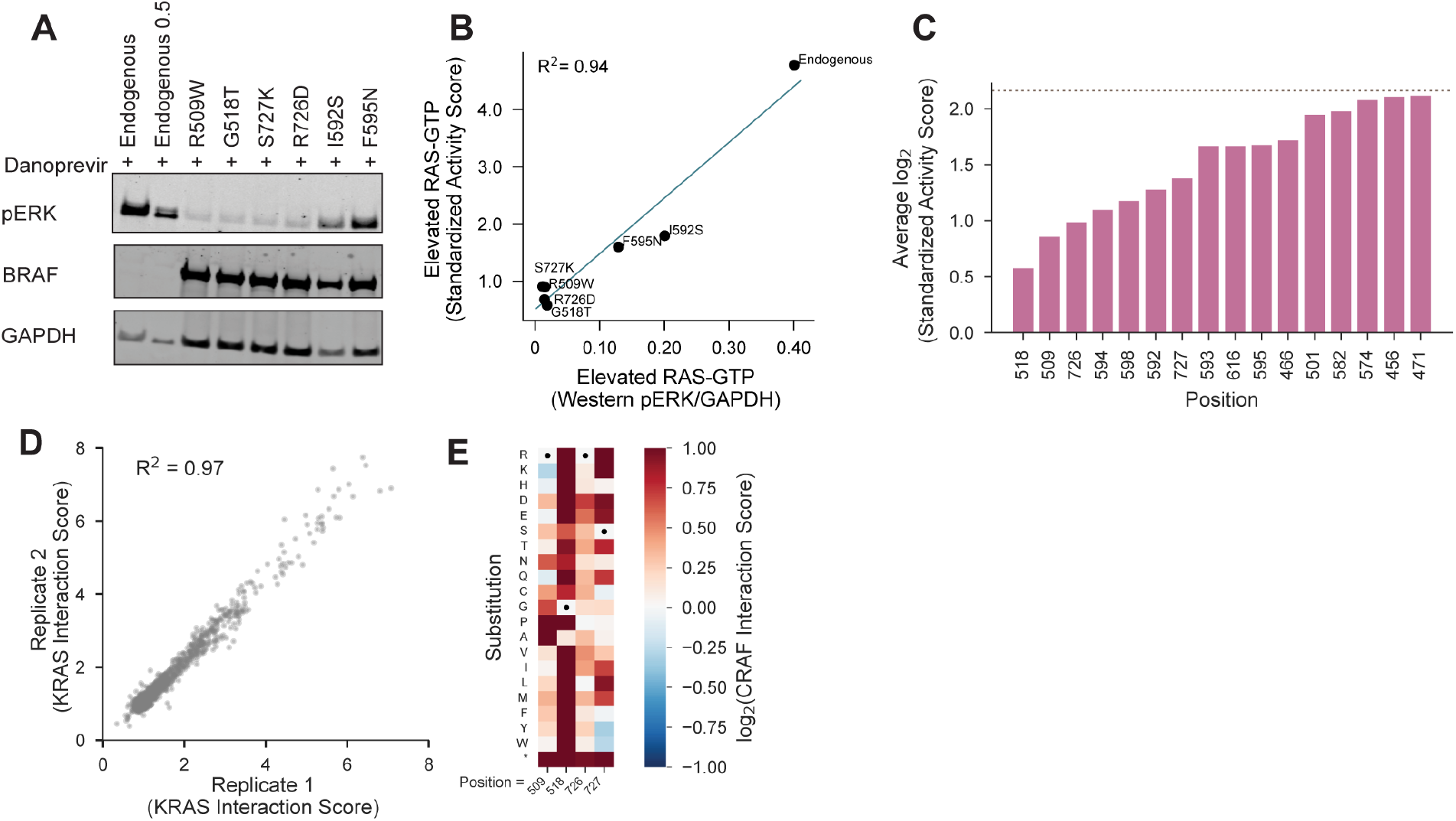
Validation of dominant negative variants, related to Figure 5. **(A)** Western blot validation of dominant negative BRAF variants under elevated RAS-GTP. Individual cell lines expressing the BRAF variants R509W, G518T, R726D, S727K, I592S, or F595N were cultured in the presence of NS3a-CIAR to elevate endogenous RAS-GTP. Lysates from each variant-expressing cell line were resolved in individual lanes by western blotting and compared with lysate from empty-vector control cells expressing NS3a-CIAR alone. Empty-vector lysate was loaded at both full and half input amounts to provide endogenous pathway-activation reference points. This blot was used to generate the quantification shown in Figure 5C. n = 2 biological replicates **(B)** Correlation between elevated RAS-GTP activity scores and pERK levels for dominant negative BRAF variants. LABEL-seq standardized activity scores measured under elevated RAS-GTP conditions were plotted against pERK/GAPDH ratios quantified by western blotting for the dominant negative BRAF variants shown in panel A. The two measurements were strongly correlated (Pearson’s R^2^=0.94). **(C)** Positional-average standardized activity scores identify BRAF sites enriched for dominant negative substitutions. BRAF positions were ranked from lowest to highest according to the mean standardized activity score across substitutions at each position. The 16 positions with the lowest positional-average scores are shown and are enriched for substitutions with dominant negative activity. The pink line indicates the endogenous pathway-activation level. At each of these 16 positions, the mean standardized activity score across all amino-acid substitutions was below the endogenous activation level. **(D)** Scatterplots comparing WT normalized KRAS interaction scores from biological replicates performed under basal Ras-GTP conditions (Pearson’s R² = 0.97). **(E)** Subset heatmap of CRAF interaction at positions 509, 518, 726, and 727. log_2_ CRAF interaction scores are shown for individual amino acid substitutions (rows) across BRAF positions (columns), with WT residues indicated as dots and a color scale from −1 to +1, centered at WT (log_2_ score = 0). Unquantified variants are shown in grey. CRAF interaction data were adapted from Simon et al.^5^

**Figure S5.**
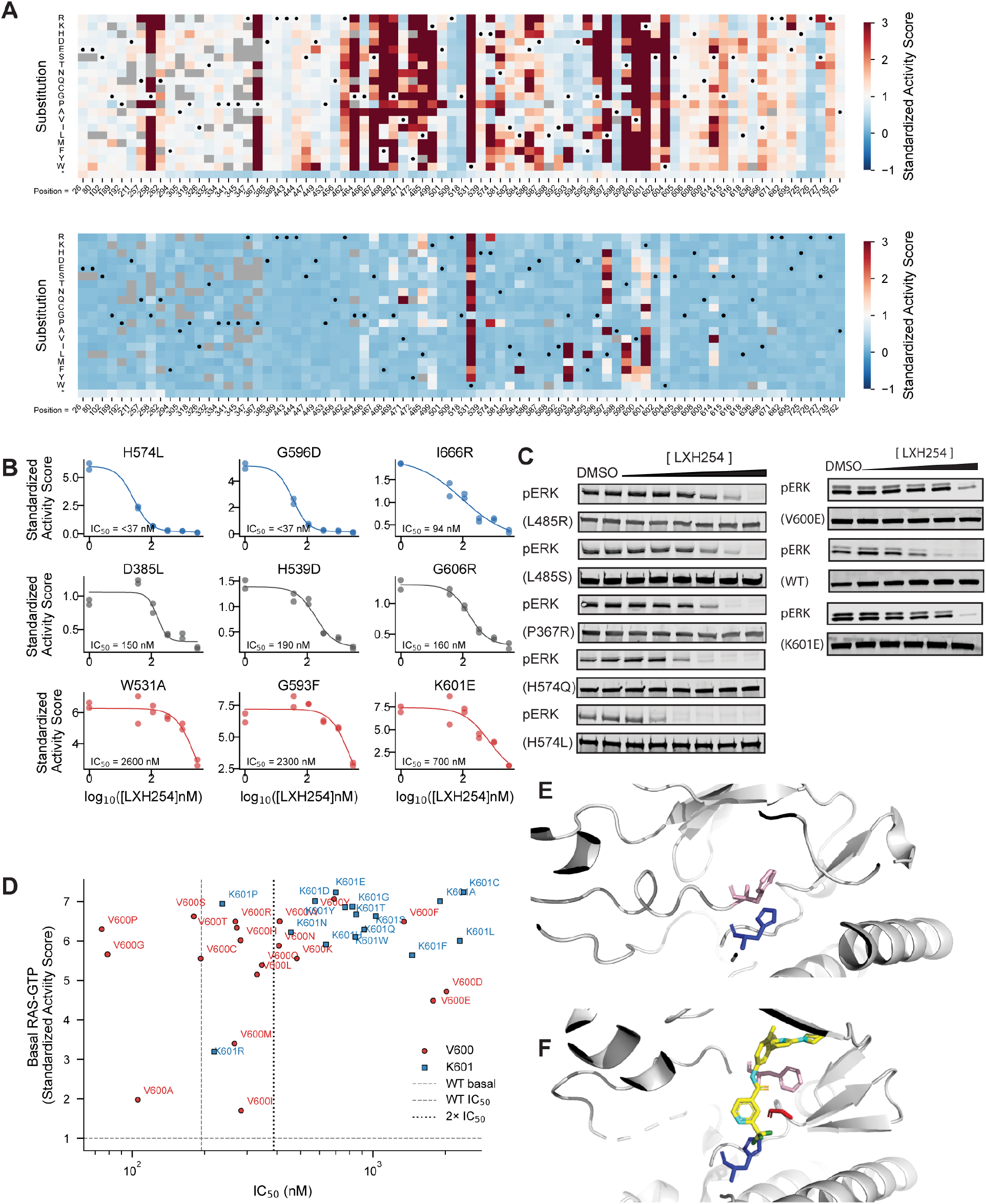
Variant dose-responses and the LXH254 binding site, related to Figure 6. **(A)** Heatmaps of log_2_ standardized pathway-activity scores measured under basal RAS-GTP conditions at the two extremes of the LXH254 titration. Cells were treated with DMSO vehicle control (0 nM LXH254; top) or 3,000 nM LXH254 (bottom). Individual amino acid substitutions (rows) are shown across the 80 profiled BRAF positions (columns); WT residues are marked with dots, and the color scale runs from −1 (inactivating) to +3 (activating). Unquantified variants are grey. (n = 2 Biological Replicates). **(B)** Representative four-parameter logistic (4PL) dose-response curves for nine BRAF variants spanning the sensitivity spectrum. Sensitizing variants (blue; H574L, G596D, I666R) exhibit IC_50_ values well below wild-type (WT IC_50_ = 190 nM); WT-like variants (grey; D385L, H539D, G606R) cluster near the WT; resistant variants (red; W531A, G593F, K601E) display markedly elevated IC_50_ values. IC_50_ values are shown for each fit. **(C)** Western blot analysis of representative BRAF variants across an LXH254 dose-response titration. Individual cell lines expressing the indicated BRAF variants were generated and treated with increasing concentrations of LXH254. Lysates from each cell line and treatment concentration were resolved in individual lanes and analyzed by western blotting for pERK. L485R, L485S, P367R, H574Q, and H574L (n = 1), V600E, WT BRAF, and K601E (n = 2). **(D)** V600 and K601 variants plotted by standardized activity scores under basal RAS-GTP (DMSO) conditions (y-axis) versus IC_50_ (x-axis). Red circles indicate V600 variants and blue squares indicate K601 variants. The horizontal dashed line denotes WT basal activity in DMSO; the dotted grey vertical line denotes WT IC_50_; the dotted black vertical line marks the resistance threshold, defined as 2-fold the WT IC_50_. Variants at these positions display diverse drug sensitivities despite similarly elevated basal activity. **(E)** Structure of autoinhibited BRAF (PDB: 6NYB), with His574 shown in blue, and Phe595 shown in pink. In this off-state conformation, His574 forms stabilizing interactions with F595. **(F)** Crystal structure of BRAF (PDB: 8F7P) bound to LXH254, shown in stick representation, with His574 shown in blue, Gly596 in red, and Phe595 in pink. In the inhibitor-bound structure, Phe595 flips outward. Loss of the His574-mediated stabilizing interaction present in the off-state (E) is expected to lower the energetic barrier to this flip, providing a structural basis for the sensitization of His574 mutants to LXH254.

## Supplementary Table Legend

- Table S2 contains scores for all enrichment regimens described in Figure 3, S1 and S2.
- Table S3 contains standardized pathway activity scores for both basal and elevated RAS-GTP used in Figure 4, 5, S3 and S4.
- Table S4 contains KRAS interaction scores used in Figure 5 and S4.
- Table S5 contains standardized pathway activity scores at all five LXH254 concentrations, along with IC50 scores, Goodness of fit is reported (R^2^) and database membership used in Figure 6 and S5.

## STAR METHODS

### KEY RESOURCES TABLE

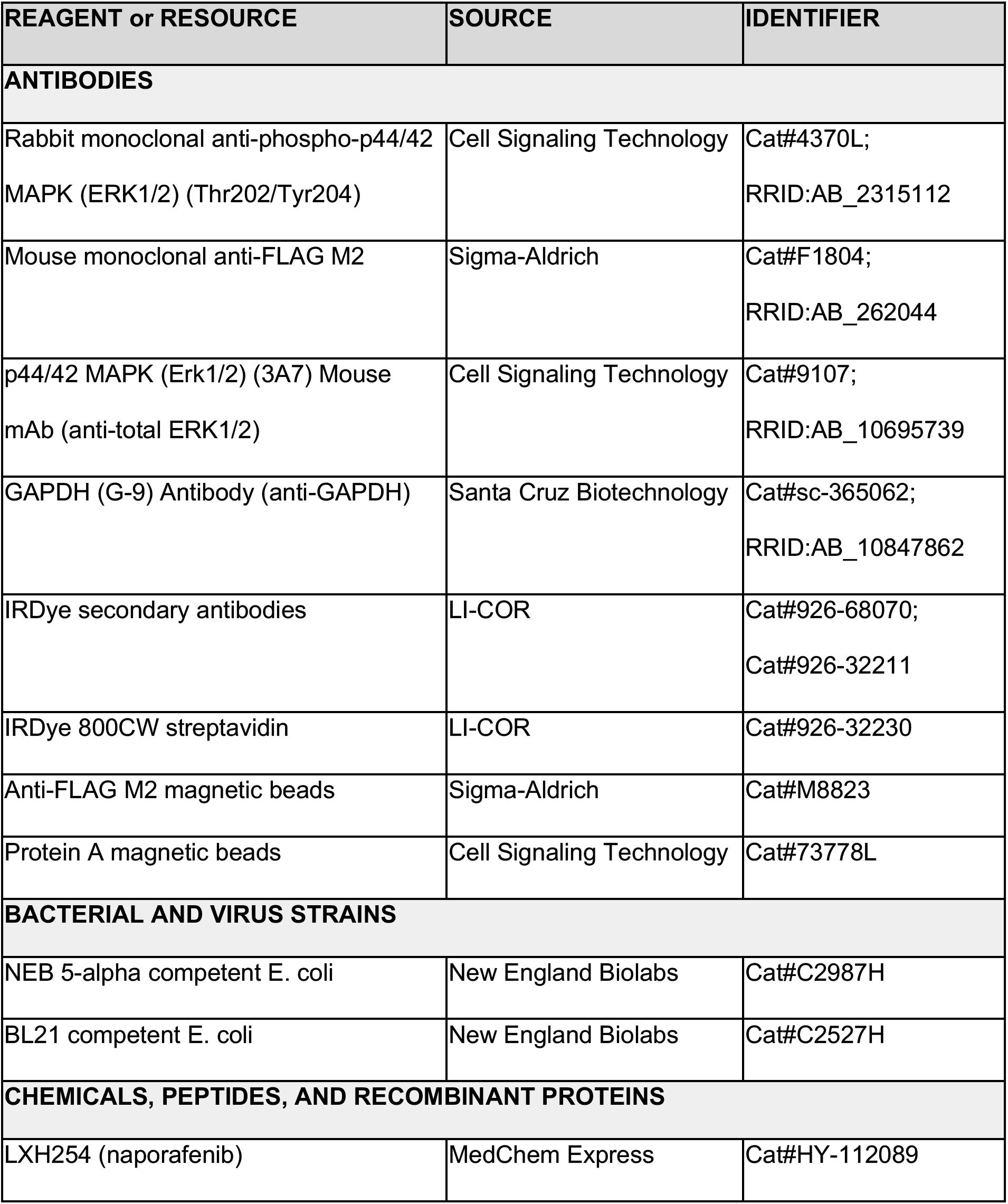
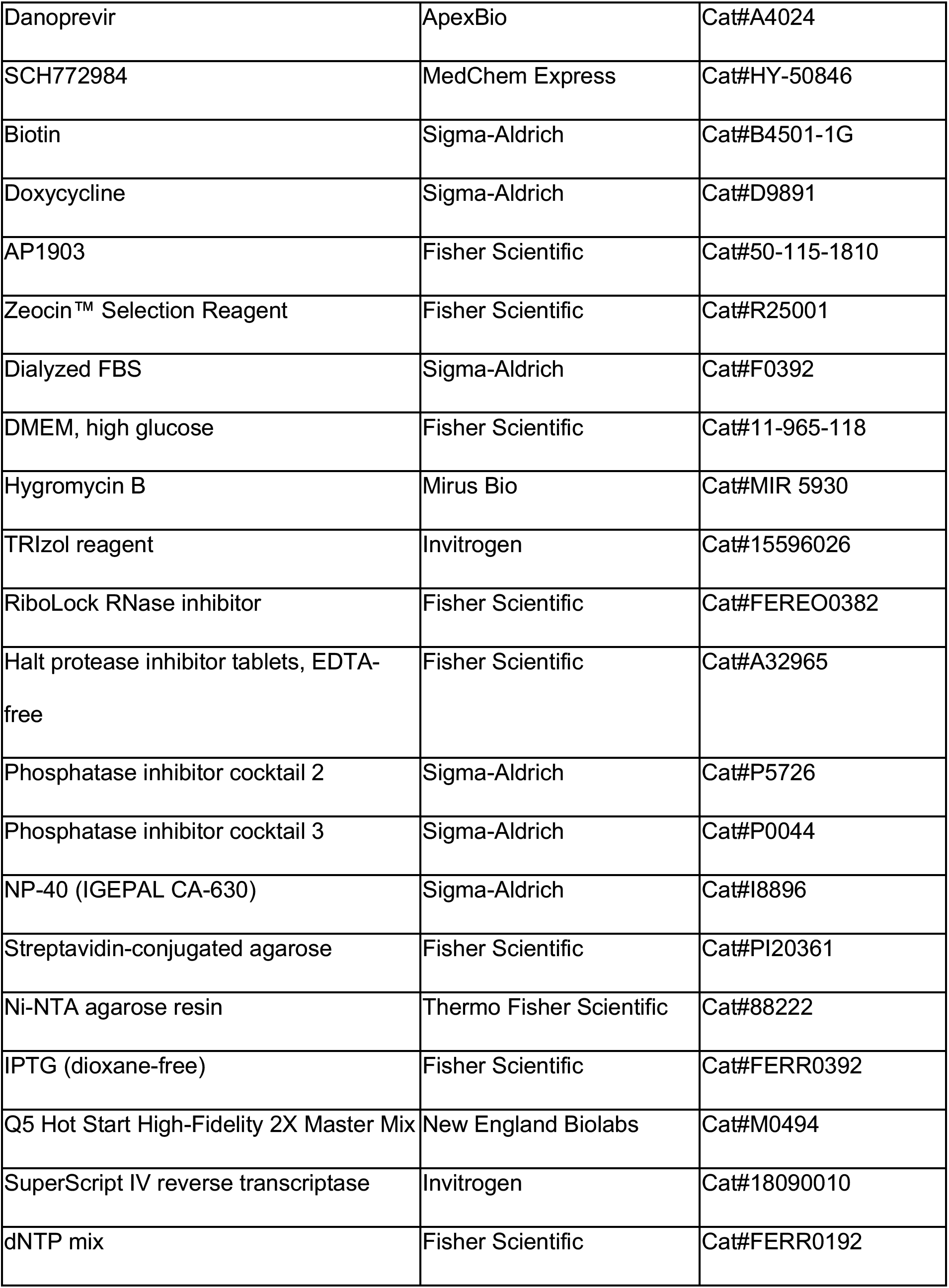
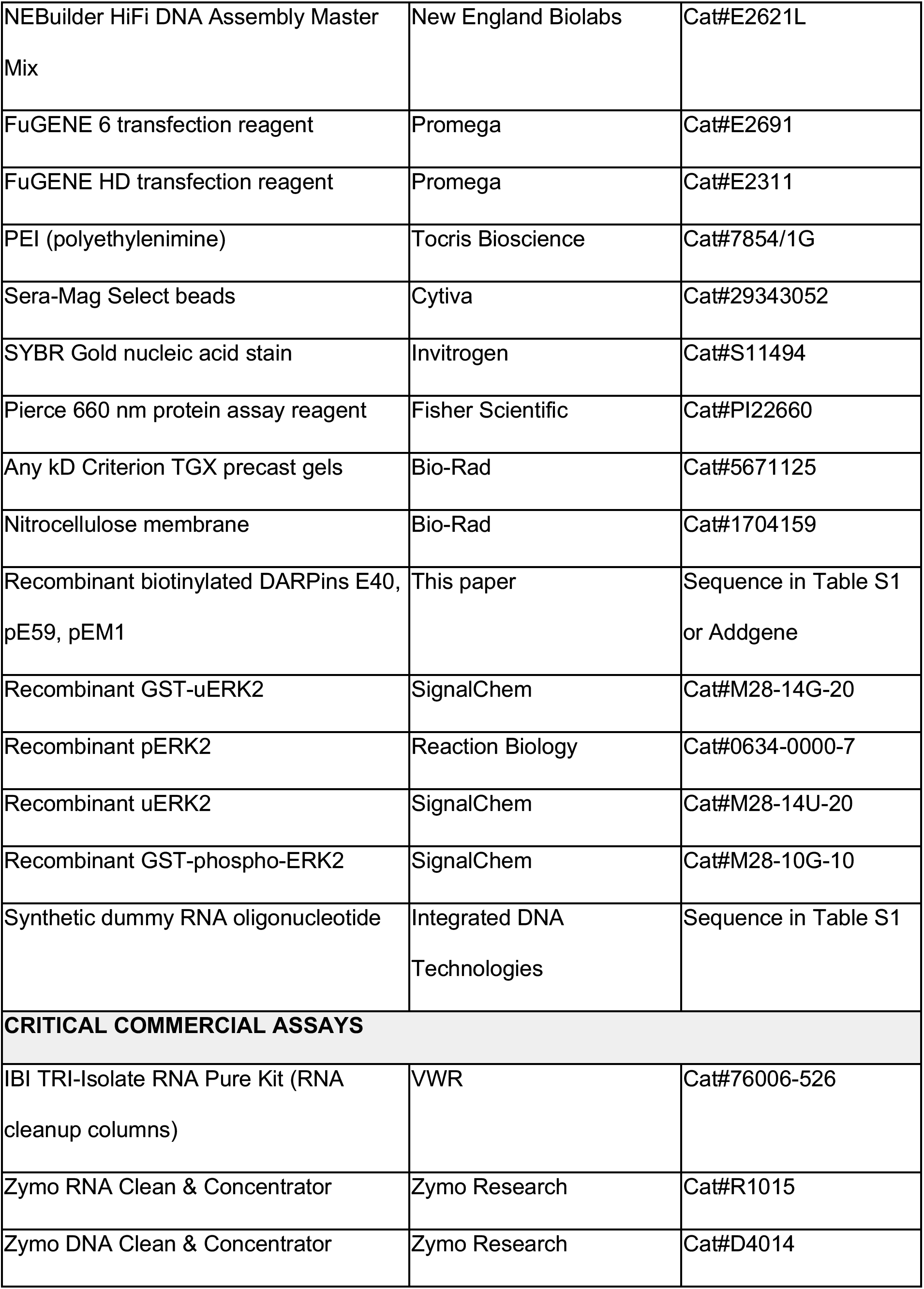
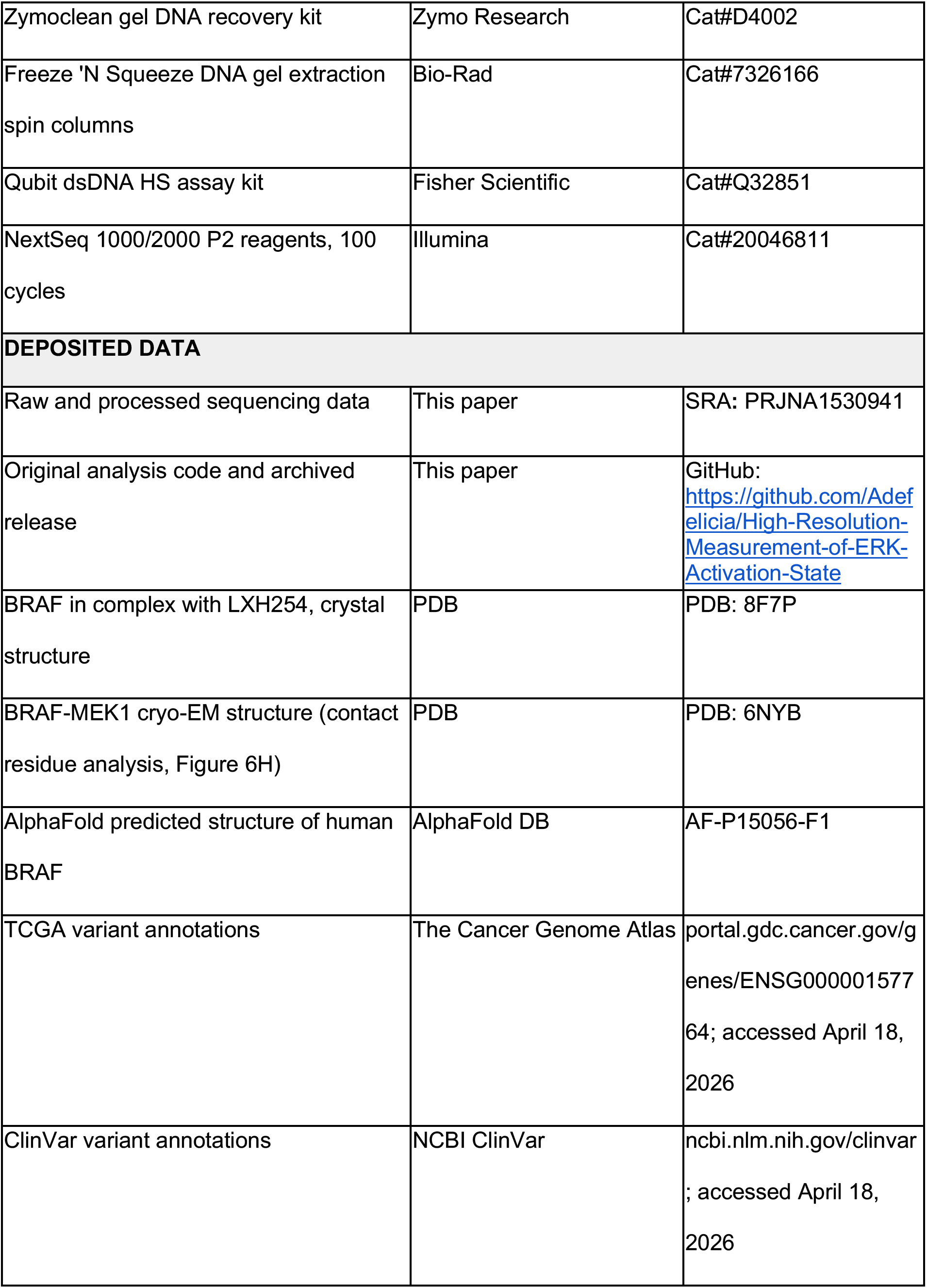
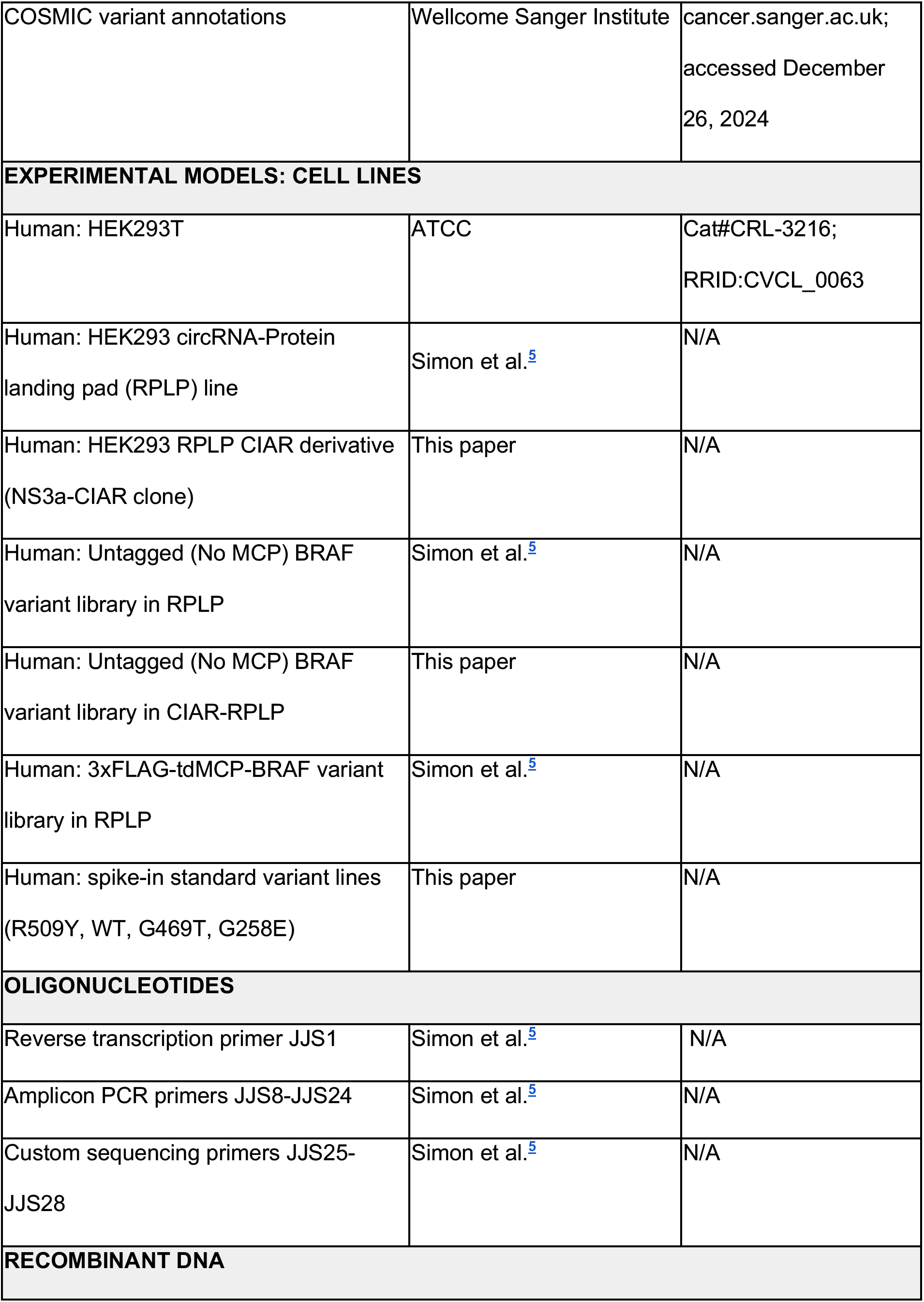
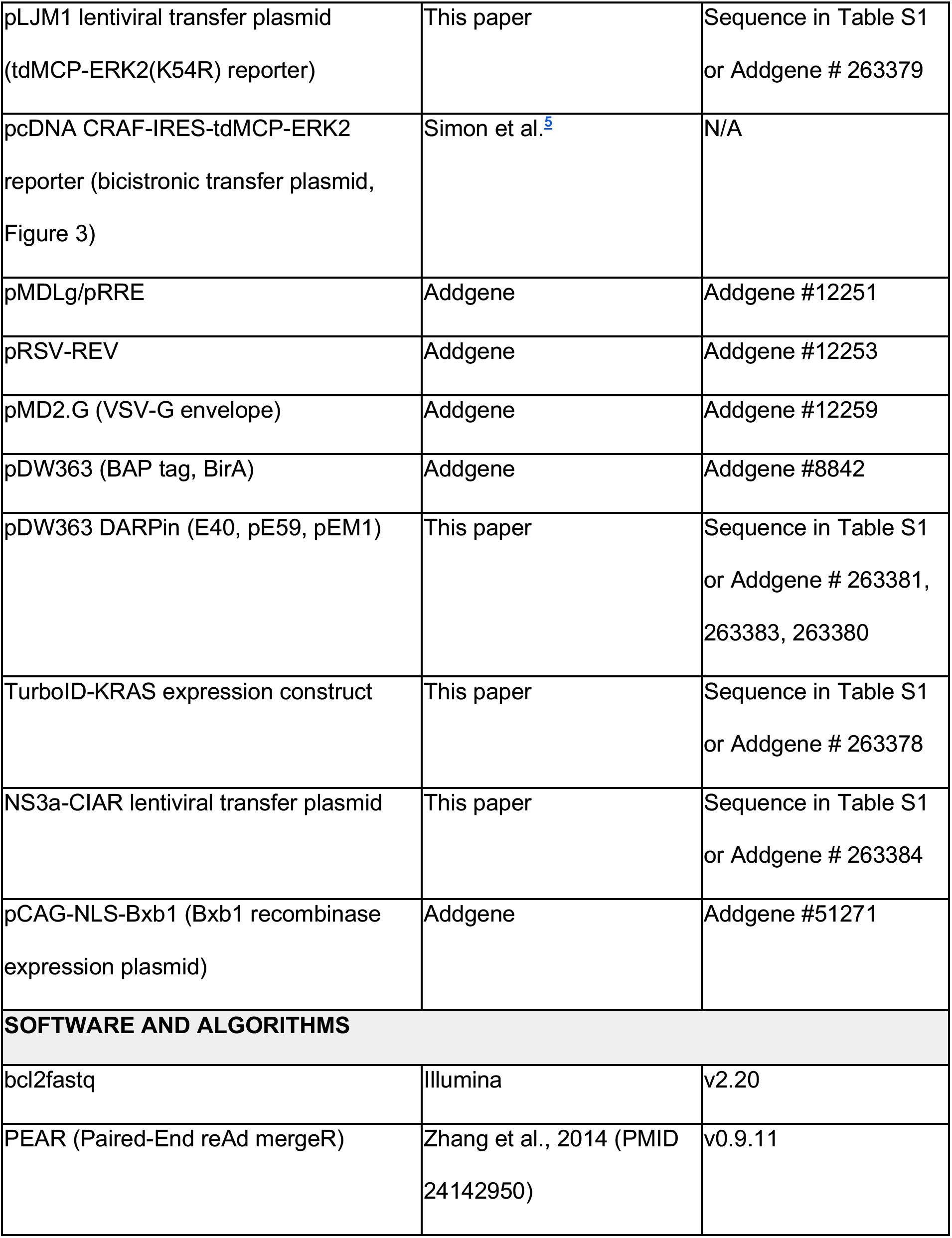
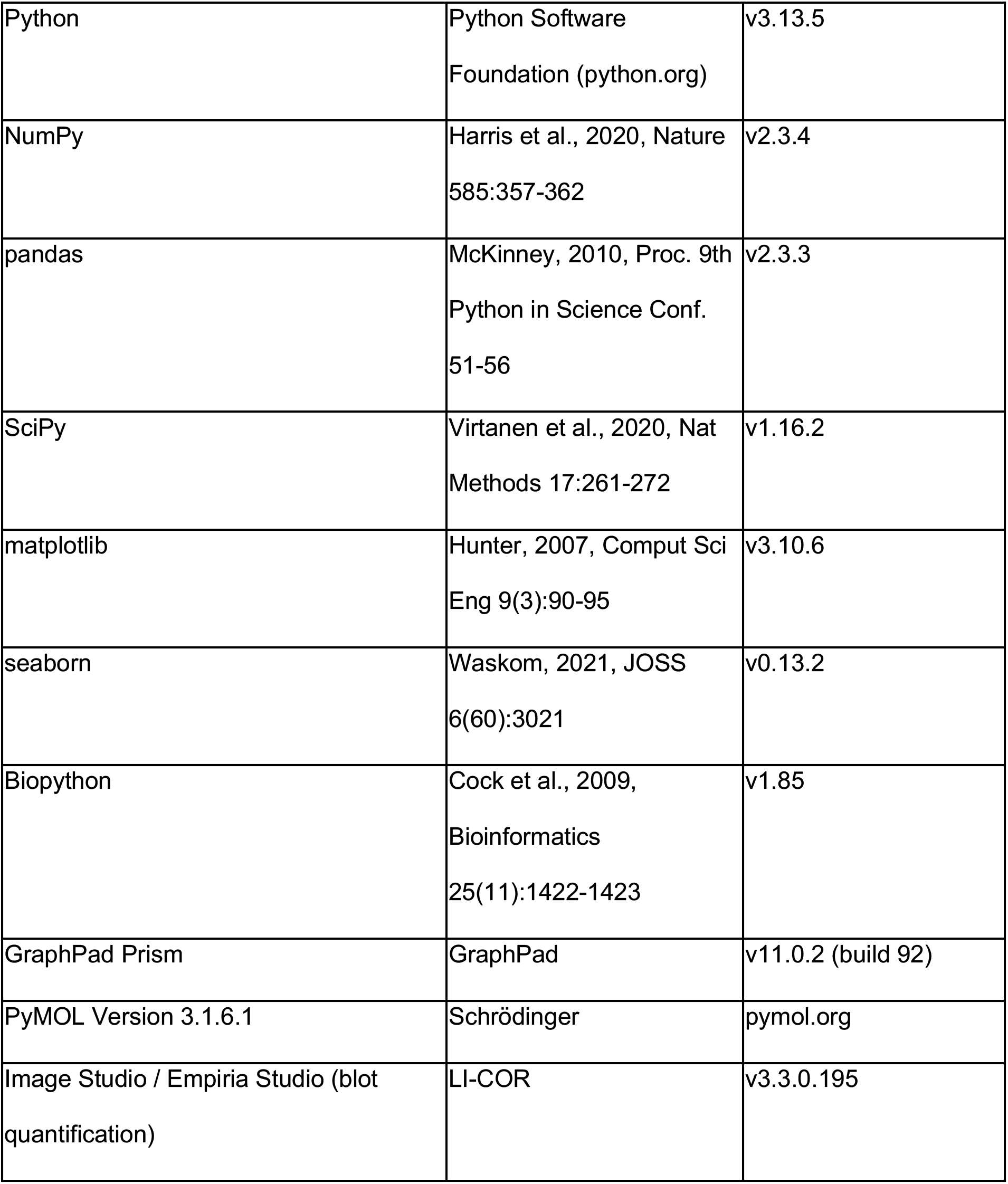

### RESOURCE AVAILABILITY

#### Lead contact

Further information and requests for resources and reagents should be directed to and will be fulfilled by the lead contact, Dustin J. Maly.

#### Materials availability

Plasmids and cell lines generated in this study are available from the lead contact upon completion of a Materials Transfer Agreement (MTA) or on Addgene.

#### Data and code availability

- Raw and processed sequencing data have been deposited at SRA [PRJNA1530941] and will be publicly available as of the date of publication. The accession number is listed in the key resources table.
- All original code has been deposited at GitHub [https://github.com/Adefelicia/High-Resolution-Measurement-of-ERK-Activation-State].
- Any additional information required to reanalyze the data reported in this paper is available from the lead contact upon request.

### EXPERIMENTAL MODEL AND STUDY PARTICIPANT DETAILS

#### Cell lines

HEK293T cells (ATCC, CRL-3216; RRID:CVCL_0063) were used exclusively for lentiviral packaging and were not used in any assay.

All pooled library experiments were performed in a HEK293-derived (Flp-In T-REx) circRNA Protein Landing Pad (RPLP) cell line, in which a single BRAF variant cassette is integrated at a defined genomic locus in each cell. PacBio barcode-to-variant maps were generated from plasmid pools of the BRAF variant library, generated by Simon et al,^5^ and were used here without modification.

For CIAR-dependent experiments (Figure 4), a clonal derivative of the RPLP line stably expressing NS3a-CIAR was generated for this study. Unrecombined RPLP cells were transduced lentivirally with a transgene cassette encoding a Zeocin-resistance gene and an mTagBFP2 fluorescent reporter. Three days post-transduction, single cells were seeded into 96-well plates by limiting dilution in growth media supplemented with doxycycline (2 µg/mL, maintained continuously as a precaution against potential transcriptional silencing of the landing-pad cassette), Zeocin (100 µg/mL), and hygromycin B (200 µg/mL); Zeocin and hygromycin B selection was maintained until cells were fully expanded. Resultant clones were expanded and screened by flow cytometry, and clones exhibiting >99% mTagBFP2 positivity were retained; a single validated clone was selected for further expansion.

The Untagged BRAF variant library was subsequently introduced into this CIAR-RPLP clone by recombination. Per 15 cm plate, a DNA mixture of 16.5 µg library plasmid, 3 µg pCAG-NLS-BxB1 (Bxb1 recombinase), and 2 µg pMAX-GFP (transfection marker) in 1 mL DMEM was combined with 80 µL FuGENE HD in 1 mL DMEM, incubated 15 min at room temperature, and added to cells. Doxycycline induction began 2 days post-transfection; AP1903 (10 nM final, from a 1,000x DMSO stock) selection began 2 days after doxycycline induction and was maintained for 24 h to eliminate unrecombined cells via the iCasp9-based negative-selection cassette.

Library cells were expanded to maintain approximately 200 cells per barcode prior to transfection or treatment.

Cells were maintained in DMEM supplemented with 10% FBS (or 10% dialyzed FBS where indicated) and doxycycline (2 µg/mL) to drive expression of the integrated variant cassette.

During expansion and prior to assay, cultures were maintained in 100 nM SCH772984 to suppress basal ERK activity; inhibitor was withdrawn 24-48 h before treatment and lysis, and all assay-day media contained no SCH772984.

#### Bacterial strains

DARPin expression constructs were propagated and expressed in E. coli BL21 (NEB, C2527H). Library and plasmid cloning used NEB 5-alpha competent E. coli.

### METHOD DETAILS

#### Plasmid construction

All plasmids were assembled by Gibson assembly of PCR products generated with Q5 Hot Start High-Fidelity 2X Master Mix DNA Polymerase and verified by whole-plasmid sequencing (Plasmidsaurus).

#### BRAF variant libraries

Both BRAF variant libraries used in this study, and the PacBio-based barcode-to-variant assignment used to construct them, were generated and described previously,^5^ no new library generation was performed for this study. Briefly, a site-saturation mutagenesis library of full-length BRAF spanning 80 selected positions was synthesized as a double-stranded DNA pool (Twist Bioscience) encoding the complete open reading frame with vector-complementary flanking sequences, cloned into a linearized attB vector by Gibson assembly, barcoded, and bottlenecked to the target number of transformants. Following recombination into the RPLP landing pad, the integrated library averaged approximately 30 barcodes per variant.

Two library formats were used: **3xFLAG-tdMCP-BRAF**, in which the tdMCP RNA-binding protein is fused to BRAF, used for the interaction assay in Figure 5; and **Untagged BRAF**, generated by excising the 3xFLAG-tdMCP module from the pre-barcoding attB library and inserting an N-terminal HA tag, used for all pathway-activity experiments (Figures 3, 4, 5, and 6).

#### tdMCP-ERK activity reporter

The reporter is a 3xFLAG-tdMCP-ERK2 fusion, with the 3xFLAG epitope at the N-terminus of the tdMCP module. Unless otherwise noted, all experiments used the kinase-dead K54R reporter. Wild-type and K54R reporters were compared directly in Figure 2; K54R was selected for all subsequent work.

The barcode is carried on a circular RNA transcribed from the integrated variant cassette and bound intracellularly by the tdMCP module, linking each recovered barcode to the BRAF variant expressed in that cell.

For Figures 2 and 3, the reporter was transiently transfected using FuGENE 6 (48 h to lysis in both cases). For Figure 2, the WT or tdMCP-ERK2(K54R) reporter was transfected into CIAR-RPLP cells (10 µg DNA per 10 cm plate; WT and K54R transfected on separate plates); cells were treated with danoprevir (1 µM, 1 hr.) or an equivalent DMSO dilution, identical to the treatment used for all other CIAR experiments, then lysed in 1 mL lysis buffer and the lysate split across the parallel- and sequential-enrichment conditions. For Figure 3, cells stably expressing the BRAF variant library were transiently transfected with a single bicistronic construct expressing WT CRAF and the tdMCP-ERK2 reporter under an IRES (17 µg DNA per 15 cm plate; one plate per replicate). For Figures 4 and 6, the reporter was delivered to library cells by lentiviral transduction.

#### Lentiviral production and transduction

Lentivirus was produced in HEK293T cells (≤ p20) plated to ∼80% confluence in two 15 cm dishes per preparation, in media containing doxycycline and 100 nM SCH772984, and transfected with 18 µg transfer plasmid (pLJM1), 9 µg pMDLg/pRRE, 4.5 µg pRSV-REV, and 4.5 µg VSV-G envelope plasmid, complexed with 108 µL PEI in 900 µL serum-free medium and incubated ∼15 min at room temperature before dropwise addition. Viral supernatant (40 mL total from two producer plates) was collected at 48 h post-transfection, filtered at 0.45 µm, and applied fresh to target cells at 7 mL per 15 cm dish. Target plates comprised one 15 cm plate of regular (untransduced) HEK293T as a transduction-efficiency control, two 15 cm plates of Untagged BRAF-RPLP (no CIAR), and two 15 cm plates of Untagged BRAF-CIAR-RPLP, with one plate corresponding to each biological replicate. Cells were expanded when confluent in media containing doxycycline and SCH772984. Medium was exchanged 48 h post-transduction to DMEM with doxycycline and FBS lacking SCH772984, and cells were assayed 72 h (3 days) after viral introduction. Transduction efficiency was assessed by flow cytometry against untransduced HEK293T.

#### Immunoblotting

Samples were resolved by SDS-PAGE on Any kD Criterion TGX precast gels (Bio-Rad) and transferred to nitrocellulose membrane. Membranes were blocked in 5% dry milk and probed with rabbit anti-phospho-ERK1/2 (Cell Signaling Technology 4370L, 1:2000), mouse anti-FLAG M2 (Sigma F1804, 1:10,000), anti-total ERK (Cell Signaling Technology 9107, 1:1000), and anti-GAPDH (Santa Cruz sc-365062, 1:5000), followed by IRDye secondary antibodies (1:10,000). Detection used a LI-COR Odyssey M and quantification was performed in Empiria Studio.

#### DARPin expression and purification

DARPin coding sequences for E40 (PDB 3ZU7), pE59 (PDB 3ZUV) and pEM1 were synthesized as eBlocks (IDT) with an N-terminal His tag and cloned into the pDW363 backbone (Addgene #8842), which supplies a BAP (Avi) tag and BirA for in vivo biotinylation.

Constructs were transformed into E. coli BL21. A single colony was grown overnight in 30 mL LB at 37°C, expanded into 1 L LB containing 0.1 mg/mL ampicillin, and grown at 37°C to OD₆₀₀ ∼ 0.6. Expression was induced with 0.5 mM IPTG in the presence of 12.2 mg/L biotin and continued 4-5 h before pelleting and storage at −80°C.

Pellets were lysed by sonication (2-3 cycles, 30 sec each, 70% output, on ice, with ∼1 min rest between cycles) in 25 mM Tris pH 8.0, 150 mM NaCl, 5 mM imidazole, 5% glycerol, 2 mM BME, and EDTA-free protease inhibitor (Pierce Mini Tablets). Lysate was cleared at 15,000 × g for 45 min and the soluble fraction bound to Ni-NTA agarose (Thermo Fisher 88222) by gravity flow.

Resin was washed once with high-salt buffer (25 mM Tris pH 8.0, 1 M NaCl, 5 mM imidazole, 5% glycerol) and twice with wash buffer (25 mM Tris pH 8.0, 150 mM NaCl, 15 mM imidazole, 5% glycerol). Protein was eluted in 25 mM Tris pH 8.0, 150 mM NaCl, 250 mM imidazole, 10% glycerol, dialyzed overnight against 20 mM Tris pH 8.0, 150 mM NaCl, 10% glycerol, 2 mM DTT, aliquoted, and stored at −80°C. Concentration of biotinylated DARPin was determined against a BSA standard curve on a Coomassie-stained gel.

#### DARPin-bead conjugation

Biotinylated DARPins were immobilized on streptavidin-coated beads (Thermo Fisher PI20361) overnight (15-18 h) at 4°C with end-over-end rotation.

For uERK DARPins, 12.8, 6.4, 3.2, or 1.6 µL of 31.2 µM E40 was combined with 10 µL bead slurry in 50 µL TBS, yielding final concentrations of 2, 1, 0.5, and 0.25 µM respectively in 200 µL lysate. For pERK DARPins, 32.65, 16.33, 8.16, or 4.08 µL of 49 µM pE59 or pEM1 was combined with 10 µL bead slurry in 50 µL 1× TBS, yielding 8, 4, 2, and 1 µM respectively in 200 µL lysate. Beads were washed twice with 10 volumes of 1× TBS before addition to lysate.

For BRAF variant library experiments, streptavidin beads were washed and incubated overnight with the appropriate biotinylated DARPins in TBS to achieve desired final concentration in lysate: (4 µM final E40 in lysate) or (2 µM final pE59 in lysate) or (0.5 µM final pEM1 in lysate).

#### Predicted binding selectivity

Fractional occupancy of purified uERK and pERK by each DARPin was calculated for ERK using published K_D_ values of DARPins^13,14^:

E40: uERK2: 6.6e-9 M, pERK2: 1.2e-6 M. pE59: uERK2: 3.5e-6 M, pERK2: 453e-9 M. pEM1: uERK2: 3.5e-6 M, pERK2: 87.1e-9 M.

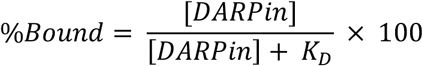

Fractional occupancy is evaluated separately for uERK and pERK for E40, pE59, and pEM1.

#### Purified protein binding assay

GST-tagged and untagged ERK species were combined in 0.8 mg/mL bacterial lysate at a final concentration of 10 nM each: GST-pERK with untagged uERK for pERK-DARPin assays, and GST-uERK with untagged pERK for uERK-DARPin assays. Parallel enrichment was performed for 1 hr. following a 3x bead washing step. Beads were eluted in 30 µL 3×SDS buffer, and boiled at 95°C for 5 min.

#### Lysis

Buffers were prepared from inhibitor- and NP-40-free modRIPA buffer (50 mM Tris, pH 7.6, 150 mM NaCl, 4 mM MgCl₂, and 5% glycerol) and supplemented with one Halt protease inhibitor tablet per 50 mL as indicated below.

● **Resuspension modRIPA:** 10 mM NaF, 2 mM PMSF, phosphatase inhibitor cocktails 2 and 3 (1:100 each), 4 µL/mL RiboLock RNase inhibitor, and 300 nM synthetic dummy RNA.
● **Lysis modRIPA:** 10 mM NaF, 2 mM PMSF, phosphatase inhibitor cocktails 2 and 3 (1:100 each), 4 µL/mL RiboLock RNase inhibitor, and 0.2% IGEPAL CA-630.
● **Wash modRIPA:** 0.1% IGEPAL CA-630.

Cells were washed with 3 mL DPBS, trypsinized with 3 mL trypsin, resuspended in 6 mL DMEM, combined with 1 mL standard spike-in cell suspension, and pelleted at 800 × g for 3 min, followed by a 1 mL DPBS wash. Pellets from a single 15 cm dish were resuspended in 0.9 mL cold resuspension modRIPA and thoroughly triturated before addition of 0.9 mL cold lysis modRIPA; lysates were rotated end-over-end for 10 min at 4°C and cleared at 17,000 × g for 10 min at 4°C.

#### Parallel enrichment

Lysate was split and incubated with uERK- and pERK-DARPin beads (or anti-FLAG and anti-pERK antibody beads) in separate reactions for 45 min at 4°C with end-over-end rotation except anti-pERK antibody beads which were incubated for 3 h, followed by three sequential washes with 10 bead volumes of cold wash modRIPA buffer.

#### Sequential enrichment

uERK was first depleted from the lysate by incubation with E40 beads (4 µM final in lysate) for 45 min at 4°C with end-over-end rotation. The flow-through (1600 µL) was transferred to pEM1 beads (0.5 µM final in lysate) - or pE59, or anti-pERK antibody beads where indicated. pE59 and pEM1 beads were incubated a further 45 min at 4°C while anti-pERK antibody beads were incubated for 3 h Beads from both steps were washed three times with 1 mL cold modRIPA with 5 min rotation per wash, then resuspended in 75 µL water plus 300 µL TRIzol and stored at − 20°C.

For Figure 5, lysate was split into 400 µL for anti-FLAG enrichment and 1400 µL for streptavidin enrichment, each for 1 hr. at 4°C. Raw streptavidin/FLAG barcode-count ratios were normalized to the wild-type ratio within each replicate - the FLAG fraction captures the total (phosphorylated plus unphosphorylated) reporter pool, while only a biotinylated, TurboID-proximal fraction is recovered by streptavidin. Beads were washed 3 × 1 mL cold modRIPA and stored as above.

#### Spike-in calibration standards

Standard variant cells, transiently expressing tdMCP-ERK2 reporters, were maintained as separate clonal populations carrying orthogonal barcodes not represented in the library.

Standards were cultured in parallel under vehicle conditions and were not exposed to danoprevir or LXH254, so that their reference values remain treatment-independent. Standards were trypsinized separately, resuspended to 2 × 10⁵ cells/mL. 1 mL of standard cell solution was added per 20 million library cells before pelleting and lysis. Each replicate received its own independent standard spike-in.

Pathway-activity standards and their assigned scores were: Std1: BRAF R509Y (0.64), Std2: BRAF wild type (1.00), Std3: BRAF G469T (4.05), and Std4: BRAF G258E (5.62); reference scores were fixed once from the basal RAS-GTP activity measurement and used to place all subsequent datasets onto a common scale.

#### Chemically inducible RAS activation (CIAR) - Figure 4

The basal and elevated RAS-GTP conditions in Figure 4 were generated using two distinct cell backgrounds: the basal RAS-GTP condition used the plain Untagged BRAF-RPLP library (no CIAR construct present) treated with an equivalent dilution of DMSO as a matched vehicle control, while the elevated RAS-GTP condition used the Untagged BRAF-CIAR-RPLP library treated with danoprevir at a final concentration of 1 µM. Both conditions were treated for 1 h in medium containing dialyzed FBS and no SCH772984, then harvested immediately. Figure 4 experiments were performed without exogenous CRAF, with the ERK reporter delivered by lentiviral transduction as described above.

#### Endogenous pathway activity level

The endogenous level was determined independently of the pooled measurement it calibrates. Clonal NS3a-CIAR lines expressing wild-type BRAF, Std2 (WT) and Std4 (G258E) - all three also present in the Figure 4 pooled experiment - were lysed alongside an NS3a-CIAR line carrying no exogenous BRAF, under the same danoprevir treatment. pERK/ERK signal was quantified by western blot for all four lines, and the three BRAF-expressing lines were regressed against their Figure 4 standard-adjusted LABEL-seq scores (Figure S3B). The pERK/ERK signal of the line lacking exogenous BRAF was then interpolated onto that regression, giving an endogenous standard-adjusted activity of 4.48 (log_2_ = 2.16), against 5.88 (log_2_ = 2.56) for wild- type BRAF under elevated RAS-GTP. This value is the dotted reference line in Figures 4B and 5A and the center of the dominant negative threshold.

#### Individual variant validation lines

Individual clonal cell lines stably expressing single BRAF variants, used for the western blot validation experiments in Figures 5B-C and S4A (dominant negative variants) and Figures 6E and S5C (LXH254 dose-response, representative variants), were generated by recombining single-variant constructs into the RPLP landing pad by the same Bxb1-mediated cassette exchange used for pooled library generation, performed individually for each variant of interest rather than as a pooled population. The individual validation lines used for the western-versus- LABEL-seq comparison in Figure 3D (WT, R682W, R682Y, H574Q, L485R, P367R, V600E) were also generated by this method and were previously published in Simon et al.^5^ Western-derived IC_50_ values (Figure 6E) were calculated by fitting a four-parameter logistic model in GraphPad Prism to Empiria Studio-quantified band intensities across the same five-point LXH254 dilution series used for the pooled library.

#### LXH254 treatment - Figure 6

Library cells (Untagged BRAF library recombined into the plain RPLP background, without CIAR) were treated for 4 h with LXH254 across a five-point, three-fold dilution series (37, 111, 333, 1000, and 3000 nM), with DMSO as vehicle. Two plates per concentration were treated, harvested, lysed, enriched, and sequenced in parallel as biological replicates. Both replicates derived from a common transduced population. Western blot validation of representative variants (Figure S5C) used the individual variant validation lines described above, treated under the same dilution series.

#### Proximity labeling - Figure 5

Library cells co-expressing 3xFLAG-tdMCP-BRAF variants and lentivirally transduced TurboID-KRAS were labeled by addition of 200 µM biotin for 15 min. Cells were harvested, lysed, enriched, and sequenced in parallel as biological replicates. CRAF interaction scores (Figure 5H) were not generated for this study; they are reused from the equivalent TurboID-CRAF proximity-labeling dataset previously published.^5^

#### Barcode amplicon preparation and sequencing

Amplicon preparation was performed as described previously,^5^ and is summarized here. RNA was extracted from enrichment beads in TRIzol, purified, and reverse transcribed with SuperScript IV and the gene-specific primer JJS1. Two successive rounds of PCR with primers JJS8-24 selectively amplified the circular RNA species and appended Illumina adapters; total cycle number across both rounds (round 1 amplification plus round 2 qPCR) ranged from 15-20 and was adjusted per sample by targeting an RFU of approximately 2000-3000 at PCR2. Reactions were pooled and cleaned prior to sequencing.

Libraries were sequenced on a NextSeq 2000 (P2, 100 cycles), yielding approximately 20 million reads per replicate. Sixteen-nucleotide barcodes were read twice using custom paired- end primers JJS25 and JJS26, and eight-nucleotide dual indices were read with JJS27 and JJS28. Base calls were converted to FASTQ format and demultiplexed with bcl2fastq (v2.20), and forward and reverse barcode reads were paired using PEAR (v0.9.11). Unique barcodes were counted, and a minimum read-count filter was determined empirically for each library by plotting barcode counts as a histogram: the cutoff (10 reads; see Filtering, below) was set at the inflection point separating higher-count sequences, representing real barcodes, from a lower- count population representing PCR and sequencing noise. Oligonucleotides used for amplification and sequencing were described previously in Simon et al. ^5^.

### QUANTIFICATION AND STATISTICAL ANALYSIS

#### Score calculation

WT Normalized parallel enrichment score (Figure 3, normalized pERK):

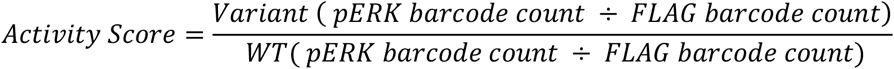

WT Normalized sequential enrichment score (Figures 3E-G):

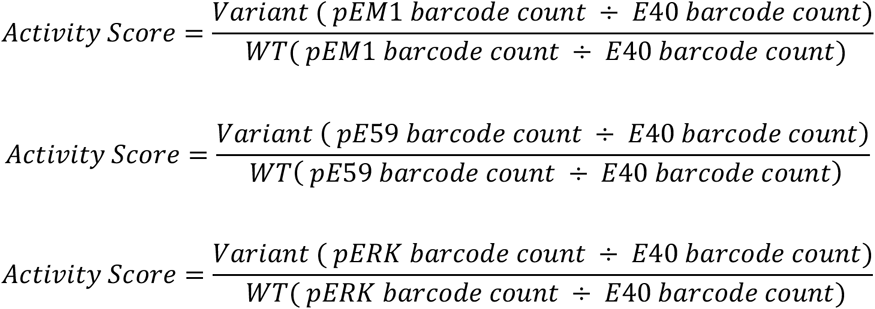

Standard-curve normalization (Figures 4, 5 and 6):

To enable quantitative comparison across libraries, assays, and batches, per-replicate scores were calibrated against the spike-in standards. For each treatment and replicate independently, an ordinary least-squares linear regression was fitted between the measured activity scores and their assigned reference scores, and all variant scores were transformed as (activity score - intercept) / slope. Per-replicate standardized scores were then averaged across the two replicates to yield the final standard-adjusted score for each variant.

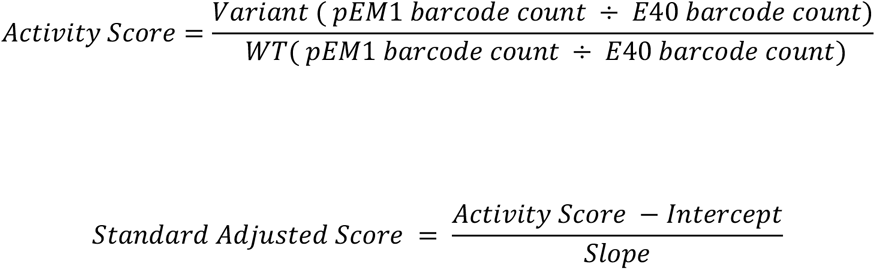

Interaction Score

For Figure 5, Interaction scores are normalized to WT.

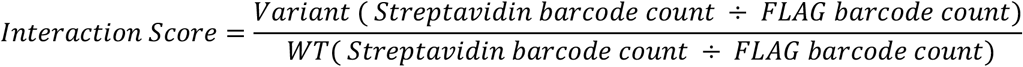

#### Filtering

To reduce undue noise, barcodes with fewer than 10 reads were excluded from analysis. Following recombination, the integrated library averaged approximately 30 barcodes per variant; after read-count filtering, variants were retained for downstream analysis only if supported by at least 4 quantified barcodes, and variants falling below this threshold were dropped. For each variant and treatment, each barcode activity score that deviated from the variant median by more than 2.5 σ within either replicate was treated as missing. This filtering step removes outlier barcodes before calculating a mean barcode score for each variant which was used as the variant activity score.

#### Functional classification

The wild-type functional zone was defined from the distribution of synonymous wild-type controls as WT ± 2σ(synonymous), with σ calculated separately per condition using that condition’s matched synonymous distribution. Variants above this zone were classified as activating, within it as WT-like, and below it as inactivating. Dominant negative variants were defined as those with an elevated RAS-GTP activity score below Endo - 2σ(synonymous), where Endo = 4.48 is the endogenous elevated RAS-GTP pathway activity level defined above.

#### IC50 determination

Loss-of-function variants were excluded prior to fitting, defined as a DMSO (0 nM) mean score below WT - 2σ(synonymous). These variants have low activity, hence no interpretable IC_50_ can be derived. Nonsense variants and standards were likewise excluded.

For each remaining variant, a four-parameter logistic model was fitted to replicate-level scores across the five concentrations and DMSO by non-linear least squares (scipy.optimize.curve_fit):

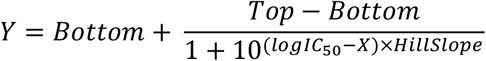

where X is log₁₀(concentration in nM). The vehicle condition was assigned a nominal concentration of 1 nM (log₁₀ = 0) for fitting.

To ensure robust fitting, curve parameters were constrained to biologically plausible values. Parameter bounds were Top [0, 10], Bottom [0, ∞), LogIC_50_ [1.0, 4.0], and HillSlope [-5, +5]; initial parameter estimates were [5, 0.5, 2.5, −1].

The upper LogIC_50_ bound of 4.0 corresponds to 10,000 nM, well beyond the maximum tested dose of 3000 nM. Fits exceeding the tested range were capped at 3000 nM for heatmap and summary visualizations and reported as ‘>3000 nM’ in dose-response plots.

The lower LogIC_50_ bound of 1.0 corresponds to 10 nM, well below the minimum tested dose of 37 nM. Fits lesser than the tested range were capped at 37 nM for heatmap and summary visualizations and reported as ‘<37 nM’ in dose-response plots.

The wild-type reference IC_50_ was 190 nM (log₁₀ = 2.28); variants were called resistant if IC_50_ was more than 2-fold higher than WT and sensitizing if more than 2-fold lower.

Goodness of fit is reported as R² (coefficient of determination) and only variants with R² ≥ 0.7 were included.

#### Database Mining

Clinical and cancer-associated BRAF variant annotation data were obtained from The Cancer Genome Atlas (TCGA) through the National Cancer Institute Genomic Data Commons (GDC; <u>portal.gdc.cancer.gov</u>), ClinVar (<u>ncbi.nlm.nih.gov/clinvar</u>), and the Catalogue of Somatic Mutations in Cancer (COSMIC; <u>cancer.sanger.ac.uk</u>). GDC and ClinVar data were accessed on April 18, 2026, and COSMIC data were accessed on December 26, 2024. For the TCGA analysis, we exported a GDC TSV file containing 308 unique BRAF somatic mutation variants and compared these records with the ∼ 1,600 BRAF variants included in our library.

Each screened BRAF variant was converted from three-letter to single-letter amino-acid notation (for example, Val600Glu to V600E) to harmonize nomenclature across COSMIC, ClinVar, and GDC. COSMIC AA Mutation entries were stripped of their leading p. prefix (for example, p.V600E to V600E) and compared directly, as COSMIC uses canonical BRAF isoform 1 residue numbering, consistent with the library. ClinVar Protein change entries, which can list multiple transcript-specific protein annotations per record, were split on commas and expanded to one row per annotation before matching. GDC/TCGA protein_change entries use BRAF isoform 2 numbering; for residues ≥437, 40 was subtracted from the reported residue number to convert variants to canonical isoform 1 numbering before matching (for example, V640E to V600E).

Variants were matched to each database by exact protein-variant identity after nomenclature harmonization. A variant was classified as database-observed if its harmonized label matched a measured BRAF variant and was reported at least once in the corresponding database. No minimum recurrence threshold was applied; for TCGA, a variant was considered observed when num_gdc_ssm_affected_cases was ≥1. Of the 308 unique BRAF variants represented in the GDC/TCGA dataset, 70 were also included and measured in the BRAF variant library.

RASopathy-associated variants were identified from ClinVar records whose phenotype or condition annotations contained one or more of the following terms: “RASopathy,” “Noonan,” “LEOPARD,” “Costello,” “cardio-facio-cutaneous,” “cardiovascular,” “Legius,” or “neurofibromatosis.” Searches were performed case-insensitively after standardizing punctuation and hyphenation. Of the ClinVar-annotated BRAF variants, 39 RASopathy- associated variants were also included and measured in the BRAF variant library. Variant activity scores were then compared across these dataset-specific variant categories, as shown in Figures S2D and S2E.

#### Structural mapping and visualization

Position-averaged scores were mapped onto protein structures in PyMOL (v3.1.6.1) using a custom Python script. Three structures were used: an AlphaFold-predicted model of full-length BRAF (AF-P15056-F1; Figures 3G and 5D), a crystal structure of the BRAF dimer bound to LXH254 (PDB: 8F7P, chain B; Figure 6G and S5F), and a cryo-EM structure of the autoinhibited BRAF-MEK1complex (PDB: 6NYB, chain A; Figure 6H and S5E).

For each residue with a quantified score, color was assigned on a diverging blue-white-red scale, linearly interpolated in log_2_ or log_10_ space as appropriate to the underlying metric, with white anchored to the wild-type reference value and values clamped at the display bounds indicated in each figure legend. Where sphere size was used to encode a second variable (percentage or count of variants meeting a defined classification, e.g., dominant negative, resistant, or sensitizing), sphere radius was linearly scaled between 0.8 and 2.5 Å across the minimum and maximum values observed for that metric across all profiled positions.

#### BRAF-MEK1 interface definition

BRAF residues forming the interface with MEK1 in the autoinhibited BRAF-MEK1 complex (PDB: 6NYB) were identified in PyMOL by selecting all BRAF residues with at least one atom within 4.5 Å of any atom of MEK1 (select BRAF_MEK_interface, byres (BRAF within 4.5 of MEK)), yielding 25 interface positions (Figure 6H). Of these, six positions (614, 615, 616, 618, 666, and 671) are included among the 80 positions profiled in the BRAF variant library and were further analyzed individually (Figure 6I).

